# Behavioural, physiological, and genetic drivers of coping

**DOI:** 10.1101/2023.08.28.555090

**Authors:** Debottam Bhattacharjee, Aníta Rut Guðjónsdóttir, Paula Escriche Chova, Esmee Middelburg, Jana Jäckels, Natasja G. de Groot, Bernard Wallner, Jorg J.M. Massen, Lena S. Pflüger

## Abstract

Animals regularly experience stressful situations, ranging from predation to social stress, yet successfully deal with them on most occasions. This adaptive mechanism, coping, reduces the adverse effects of stressors through behavioural and physiological efforts, failing to which may result in reduced fitness. However, considerable variation in coping is observed. Unlike in humans, coping is often considered a personality trait in non-human animals due to construct similarity, resulting in conceptual ambiguity. Besides, limited multidisciplinary research has rendered comprehending the drivers of coping in animals challenging. We assessed repetitive behavioural coping or coping styles (*n=30*), emotional arousal (*n=12*), and consistent inter-individual differences, i.e., personalities (*n=32*) of long-tailed macaques (*Macaca fascicularis*) using observations, ecologically relevant experiments, and infrared thermography. We finally investigated the association of coping with a Valine/Methionine polymorphism encoded by the Catechol-O-methyltransferase (COMT) gene (*n=26*), which is widely known for its involvement in human stress regulation. Our findings suggest that personality and the presence of the human equivalent *COMT* Val^158^Met polymorphism in long-tailed macaques are associated with ‘emotion-focused’ and ‘problem-focused’ coping styles. These coping styles were consistent with emotional arousal as measured with infrared thermography. We discuss these proximate drivers of coping for a better understanding of its evolution in primates.

## Introduction

Human and non-human animals are repeatedly exposed to unpredictable and uncontrollable stressors ^1^. The mechanism of coping diminishes the adverse effects of stressors through behavioural and physiological efforts ^2^. Coping has adaptive significance from an evolutionary perspective, as failing to cope with stressful situations may result in reduced fitness outcomes ^3^. Substantial variations are, however, observed in how individuals cope with identical stressors, and if those variations are consistent, they are considered coping styles ^4^. Several theoretical models have been proposed and modified in the last few decades to define the key distinctive properties of coping ^5^. For example, the ‘problem versus emotion focus’ model suggests that while problem-focused coping is directed to encounter and diminish the stressor itself, emotion-focused coping finds alternate ways to minimise the harm of distress ^6^; the ‘engagement versus disengagement’ model highlights that engagement or approaching coping is primarily problem-focused but can include components of emotion-focused coping too, whereas, disengagement coping is attributed to avoidance ^7^; the ‘proactive coping model’ points out that coping can occur proactively even before any introduction of stressor, but the components are typically problem-focused ^8^. Therefore, although the proximal goals of the underlying problem- and emotion-focused coping may appear different, they are not entirely distinctive in their mechanisms and can complement each other ^9^. Despite evident theoretical and empirical advancement of research in humans, little is known about the evolution of coping. This is mainly due to conceptual incoherence of coping in non-human animals, a restricted focus on a few taxa, and limited multidisciplinary research designs. Identifying the proximate mechanisms of coping and their roles in non-human animals may thus provide significant insights into its evolution.

In order to explain the behavioural and physiological underpinnings of coping, a ‘two-tieŕ model posits that the responses to stressful situations have two independent dimensions, coping style (i.e., a qualitative behavioural dimension: ‘proactive-reactive continuum’ and reactivity of the sympathetic autonomic nervous system) and stress reactivity (i.e., a quantitative dimension: through Hypothalamic-pituitary-adrenal/HPA axis activity) ^10^. This model received support from empirical studies conducted mainly on fishes and small mammals, like rodents ^11–13^. A proactive coping style (also problem-focused) indicates an active response to stressors with a higher propensity to take risks and form rigid routines, while a reactive coping style (often emotion-focused) indicates a passive response and higher behavioural flexibility ^14^. Typically, active copers exhibit low inhibition control, high novelty seeking and frequent aggressive and impulsive behaviours ^15,16^; on the contrary, avoidant copers do not directly engage with the stressors or show aggressive and impulsive behaviours. At the physiological level, avoidant coping is associated with higher HPA axis reactivity than active coping ^14,17^. However, according to the two-tier model, avoidant copers can show a low HPA axis- or ‘emotional’ reactivity due to the independence of coping style and HPA axis (synonymously used as emotionality) reactivity. For example, ‘docile’ but not ‘shy’ individuals can have reactive coping styles yet still show low emotionality. Notably, emotionality in coping has primarily been investigated by measuring stress hormone concentrations. Nevertheless, from a neurophysiological perspective, emotional states are associated with the activities of sympathetic and parasympathetic nervous systems, which may increase and decrease blood pressure, resulting in blood flow alterations underneath the skin ^18,19^. This change in temperature, especially in the face, can thus be used as a reliable alternative for measuring emotional arousal during coping. But, limited information is available on whether such a non-invasive methodological tool can capture emotionality in coping.

A key proximate driver of variation in coping styles may be personality. Personality, i.e., consistent inter-individual differences ^20^, are similar in constructs to coping styles. While both personality and coping styles can be attributed to individual-level differences, the question remains whether coping styles are mere reflections of personalities in stressful contexts. Scientific research in humans predominantly keeps these two concepts fundamentally distinct, and their associations are studied extensively ^20–23^. Coping, unlike personalities, was found to be less stable over time with adjustments ^21,24^ and had very low heritability ^22^. Furthermore, little to no evidence suggested an overlapping genetic basis of coping styles and personalities ^22^. In contrast to the human literature, non-human animal research often considers coping styles and personalities synonymous. For instance, the ‘boldness-shynesś personality dimension ^25^ (typically used in birds, fishes, etc.) is used interchangeably with proactive-reactive coping styles ^14^. Consequently, the concepts of coping (in human and non-human animals) and its (in)dependence on personality need more clarity as variable trait characteristics are subject to selection pressure and are of immense importance.

Conventionally, research on coping has taken an integrated approach of behaviour and neurophysiology only. As a result, candidate genes of the HPA axis are yet to be significantly explored as potential causal mechanisms. Notably, both coping and personality have some genetic basis, which may work independently or be interlinked through feedback loops ^26–28^. For example, zebrafish (*Danio rerio*) lines selected for proactive and reactive coping styles show distinct basal neurotranscriptomic states ^29^, and the Swiss sublines of Roman high- and low-avoidance rats selected for good versus poor performance in exploration tasks exhibit divergent stress responses and coping styles ^30^. These findings emphasise the need for studies on key candidate genes regulating stress coupled with behavioural and physiological mechanisms. Such a comprehensive approach is essential for gaining a holistic understanding of coping.

A key candidate gene underlying varying coping styles in humans, and potentially other animals is the so-called *COMT* (Catechol-O-methyltransferase) gene. COMT is a key enzyme encoded by the *COMT* gene in catecholamine catabolism ^31^. This enzyme is responsible for the inactivation of dopamine, adrenaline and noradrenaline neurotransmitters ^32^ and is accountable for more than 60% of dopamine clearance in the brain ^33,34^. In humans, a single nucleotide polymorphism (SNP) located in the coding region of the *COMT* gene (codon 158, exon 4) has been shown of functional consequences. At this locus, a transition from guanine (G) to adenosine (A) results in changing the amino acid product from valine (Val) to methionine (Met). The resulting gene products differ significantly in thermostability and, subsequently, in COMT enzymatic activity. The Val variant is considered more stable and active than the thermolabile Met encoding product ^35^. This functional *COMT* Val^158^Met polymorphism (dbSNP: rs4680) has been studied extensively in humans. Individuals with the G/G (Val) variant are associated with ‘warrior’ phenotypes, like higher aggression ^36^ and impulsive behaviour ^37^, whereas emotional problems ^38^ and increased reactivity to stress ^39^ are found in individuals with the A/A (Met) variant. In non-human primates (NHP), *COMT* polymorphism is linked with aggression, dominance, and stress ^40,41^. To our knowledge, however, in the NHP genus *Macaca,* the human equivalent *COMT* Val/Met polymorphism was only identified in Assamese macaques, where a moderate relation between rank and aggression was found ^41^. Although crucial from a mechanistic point of view, no studies to date have investigated the association of *COMT* polymorphism and coping in non-human animals.

The current study uses a multidisciplinary framework to investigate the coping behaviour of long-tailed macaques (*Macaca fascicularis*), a group-living NHP species. We conducted ecologically relevant predator exposure experiments repeatedly (see ^42,43^) to assess coping styles. Encounters with predators can be unpredictable and uncontrollable, following the definition of a stressor ^1^, which further allows individuals to cope with the situation. An infrared thermal imaging method was employed to detect emotional arousal during predator exposure ^44^. A drop in nose temperature is typically associated with high emotional arousal in NHP ^19,45^ and thus can capture emotionality in coping. We furthermore examined personalities in the long-tailed macaques using a comprehensive multi-method ‘bottom-up’ approach of repetitive behavioural observations and experiments (cf. ^42^). Since relatively steep hierarchical structures are found in long-tailed macaque societies ^46^, we also determined the dominance-rank relationships as potential predictors of coping and its variations. Finally, we extracted DNA from blood samples of a subset of subjects to investigate the potential existence of *COMT* polymorphism in the human equivalent *COMT* Val/Met site (exon 4) of long-tailed macaques.

We hypothesised that personality and underlying *COMT* genotype would moderate behavioural coping styles and emotional arousal, explaining individual variations. Due to the nature of the ‘bottom-up’ approach, personality dimensions were not predetermined and thus, informed predictions could not be made. Yet broadly, we predicted that activity, aggression, and exploration-like traits would be positively associated with problem-focused coping styles, and neuroticism-like traits would be positively associated with emotion-focused coping styles. As high-ranked individuals are often associated with boldness in despotic societies ^47,48^, we expected them to have problem-focused coping styles compared to their subordinate counterparts. Furthermore, we expected behavioural coping styles and emotional arousal to correlate. Thus, compared to a baseline measure, individuals with problem-focused coping styles are expected to maintain higher average nose temperatures (i.e., low emotional arousal), or recover faster from an initial drop in nose temperature than individuals with emotion-focused coping styles. Finally, we expected the underlying *COMT* genotypes, if the human-like SNP is present in long-tailed macaques, to be associated with different coping styles in our test subjects. Given the higher emotional resilience attributed to the human G/G genotype (Val carriers) ^49–51^, we would expect long-tailed macaques having the *COMT* G/G genotype (Val) to be less emotionally aroused and to exhibit more problem-focused coping styles compared to macaques carrying the A/A (Met) variant.

## Results

### Behavioural coping styles

Predator models were validated as efficient stressors, as self-directed behaviours were more frequent during predator exposure (0.49 ± 0.33 per minute) than the non-predator baseline (0.27 ± 0.10 per minute) condition (Wilcoxon Signed-Rank test: z = −3.42, r = 0.66, p < 0.001, **Supplementary Fig. 1**).

Considering the behavioural variables we expected to relate to coping (**Supplementary Table 1**), two non-correlating latent factors were extracted using an exploratory factor analysis (EFA). These two factors explained 37.6% and 22.4% of the total observed variance, i.e., 60% cumulatively. The first factor included positively loaded variables of *close ground, conspecific affiliation,* and negatively loaded variable of *far* (**Fig. 1**). In contrast, the second factor had only two positively loaded variables – *predator aggression* and *conspecific aggression* (**Fig. 1**). Clearly, the second factor loaded variables relevant to a direct engagement with the stressor; thus we labelled it as ‘problem-focused coping’ (cf. ^23^). The first factor, apart from a close distance, did not indicate any direct engagement with the stressor. Instead, a reliance on conspecifics through affiliation was found. Thus, the first factor had an emotional component, and we labelled it as ‘emotion-focused coping’ (cf. ^23^). Individual scores were obtained from the two factors and plotted against each other using a scatter plot (cf. ^52^).

**Fig. 1.**
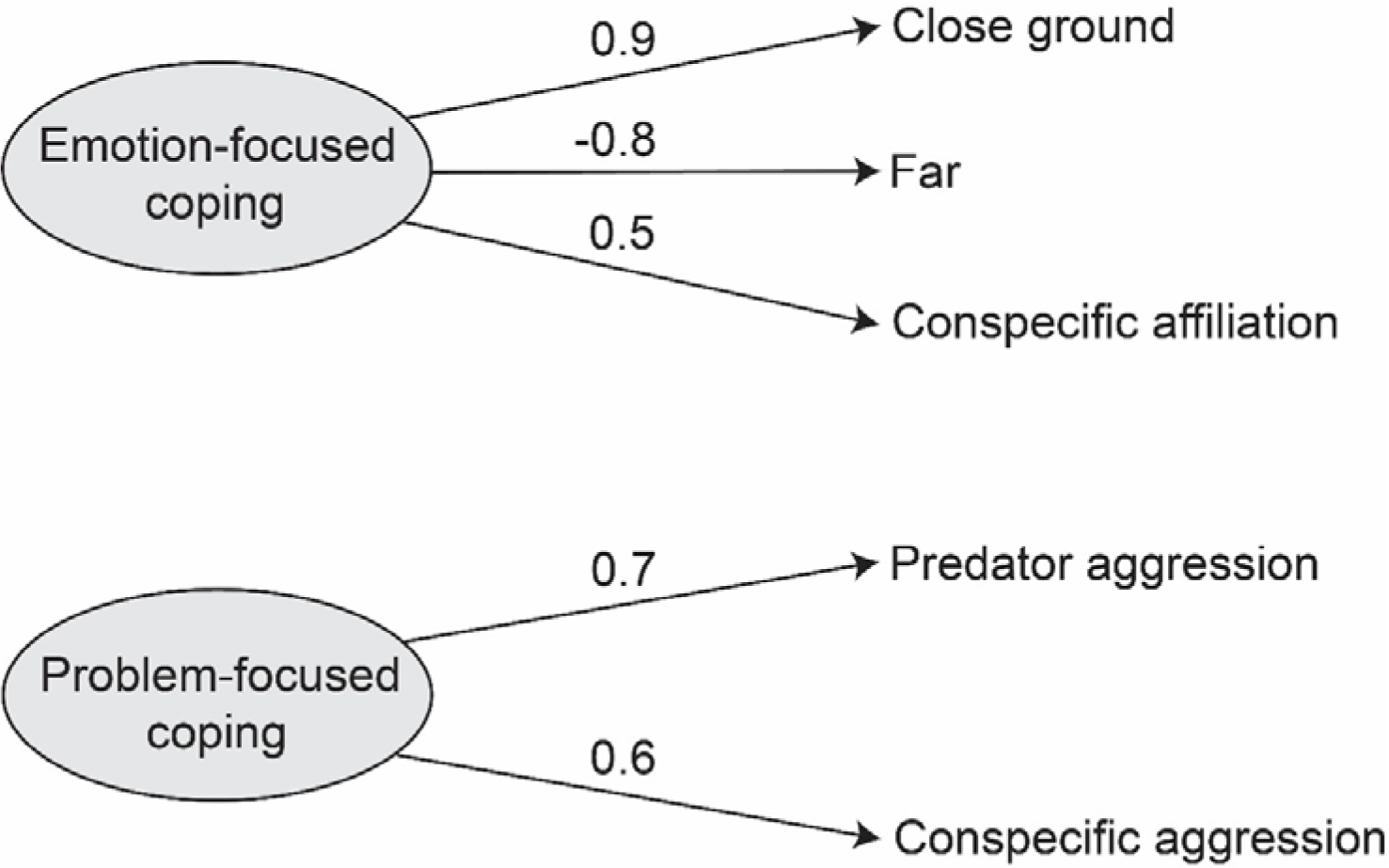
Exploratory factor analysis (EFA) showing two latent factors. Latent factors, namely emotion-focused- and problem-focused coping, were identified (n=30). The behavioural variable loadings are indicated by the positive and negative values.

The plot helped determine the general categorisation of the individuals into problem-focused, emotion-focused, mixed coping styles and individuals with low reactivities (**Supplementary Fig. 2**). Out of the 30 individuals tested, nine appeared to have primarily a problem-focused coping style, seven had primarily emotion-focused- and six had mixed coping styles. The remaining eight individuals had low reactivities, i.e., these individuals scored below the median population value for both problem-focused (median = 0.15) and emotion-focused (median = −0.11) factors and could not be assigned to a particular single coping style. The six individuals categorised into mixed coping had significantly higher absolute values for the problem-focused coping factor in comparison to their scores on the emotion-focused dimension (Wilcoxon Signed-Rank test: z = −2.15, r = 0.88, p = 0.03); therefore, it was decided to combine problem-focused- and mixed copers together (n = 15). This combined group, defined as problem-focused, along with the emotion-focused (n = 7) copers, were used for the rest of the analyses. See **Supplementary Note 1** for group-level details.

### Emotional arousal

According to the best-fitted model (**Supplementary Table 2**) on our thermography data, we found a significant effect of coping style on the nose temperature change (LME: t = −2.490, *η^2^* = 0.45, p = 0.03). In comparison to an initial 2-10 min time window (baseline), individuals with problem-focused coping styles regained and maintained higher average mid-nose temperatures (average change in temperature = 2.54 ± 3.24 °C) than those with emotion-focused coping styles (average change in temperature = 0.03 ± 1.44 °C, **Fig. 2**). The temperatures remained consistent for either coping style during the two different time windows (2-10 min vs 10-20 min - LME: t = 1.815, p = 0.08; 2-10 min vs 20-30 min - LME: t = 1.889, p = 0.07). Note that the baseline nose temperatures remained comparable between problem- and emotion-focused copers (Mann Whitney U Test: z = −1.314, p = 0.17, **Supplementary Fig. 3**). We did not find any effect of sex on emotional arousal (LME: t = 1.408, p = 0.19).

**Fig. 2.**
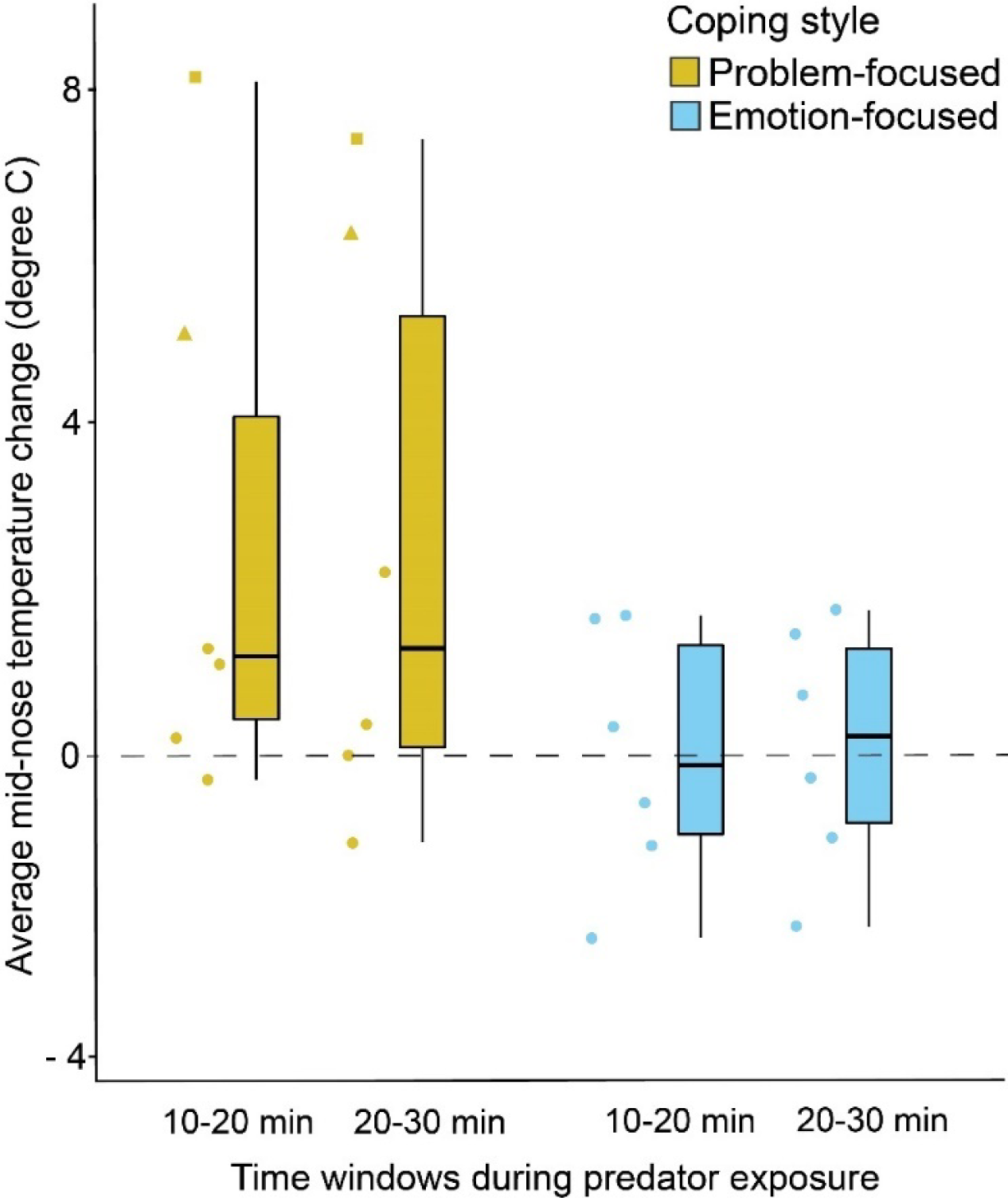
Mid-nose temperature change during predator exposure. The box plot shows the average mid-nose temperature change during predator exposure. Individual data points are represented using solid dots, squares and triangles (n=12). The squares and triangles indicate individuals with *COMT* Val/Met G/G genotype. Boxes represent interquartile ranges, and whiskers represent the upper and lower limits of the data. The horizontal bars within the boxes represent the median values.

A validation test found no significant difference between problem- and emotion-focused copers regarding general activity (problem-focused copers: 654.28 ± 253.15 s/hour; emotion-focused copers: 771.98 ± 246.38 s/hour; Wilcoxon Rank-Sum test: z = 1.106, p = 0.31), suggesting that the observed nose temperature changes were not due to general movement and foraging, rather an effect of the activities of the sympathetic autonomic nervous system, further confirming the emotional response we were aiming to measure.

### Personality

Based on the behavioural variables extracted from focal observations and novelty experiments, a principal component analysis (PCA) provided us with three PCs (with eigenvalues >1), cumulatively explaining 78% of the total observed variance (**Table 1**). We labelled the PCs ‘Activity-sociability’, ‘Affiliation’, and ‘Exploration’. The effects of age and sex on personality traits are provided in **Supplementary Note 2** and **Supplementary Tables 3** and **4**. Individual scores were obtained for the three traits (**Supplementary Table 5**).

**Table 1.** Output of the principal component analysis. Variables, factor loadings (> ±0.5), attribute communalities, and variance explained by each principal component are provided in the table.

| Variables | Activity-Sociability | Affiliation | Exploration | % communalities |
| --- | --- | --- | --- | --- |
| Approach | 0.93 |  |  | 94.14 |
| Leave | 0.92 |  |  | 72.28 |
| Follow | 0.85 |  |  | 82.26 |
| Pass by | 0.84 |  |  | 87.59 |
| Leave passive | 0.83 |  |  | 74.82 |
| Sit | -0.83 |  |  | 75.76 |
| Social play | 0.79 |  |  | 87.51 |
| Proximity |  | 0.92 |  | 80.88 |
| Groom |  | 0.75 |  | 77.41 |
| Handling container |  |  | 0.97 | 87.3 |
| Object manipulation |  |  | 0.66 | 81.57 |
| Hang |  |  | 0.54 | 84.17 |
| **% variance** | **46** | **17** | **15** |  |

### Personality, dominance hierarchy, and behavioural coping styles

We did not find any effect of the three personality traits, age and sex, on problem-focused coping (**Supplementary Table 6**). However, according to the best-fitted model (**Supplementary Table 7**), we did find an effect of the personality trait affiliation (LME: t = - 6.00, *η^2^* = 0.95, 95% CI = 0.32, 1, p = 0.02, **Fig. 3a**) and of sex (LME: t = −6.81, *η^2^* = 0.96, 95% CI = 0.42, 1, p = 0.02, **Fig. 3b**) on emotion-focused coping. An inverse association between affiliation and emotion-focused coping was found, while females (n = 5; 1.41 ± 0.92) had higher emotion-focused coping scores than males (n = 2; 0.45 ± 0.21). We did not find any effect of the personality trait exploration on emotion-focused coping (LME: t = 2.12, p = 0.16) but found a non-significant negative trend of age (LME: t = −3.94, p = 0.05). Note that the non-significant activity-sociability fixed effect was dropped from the best-fitted model during the model selection process.

**Fig. 3.**
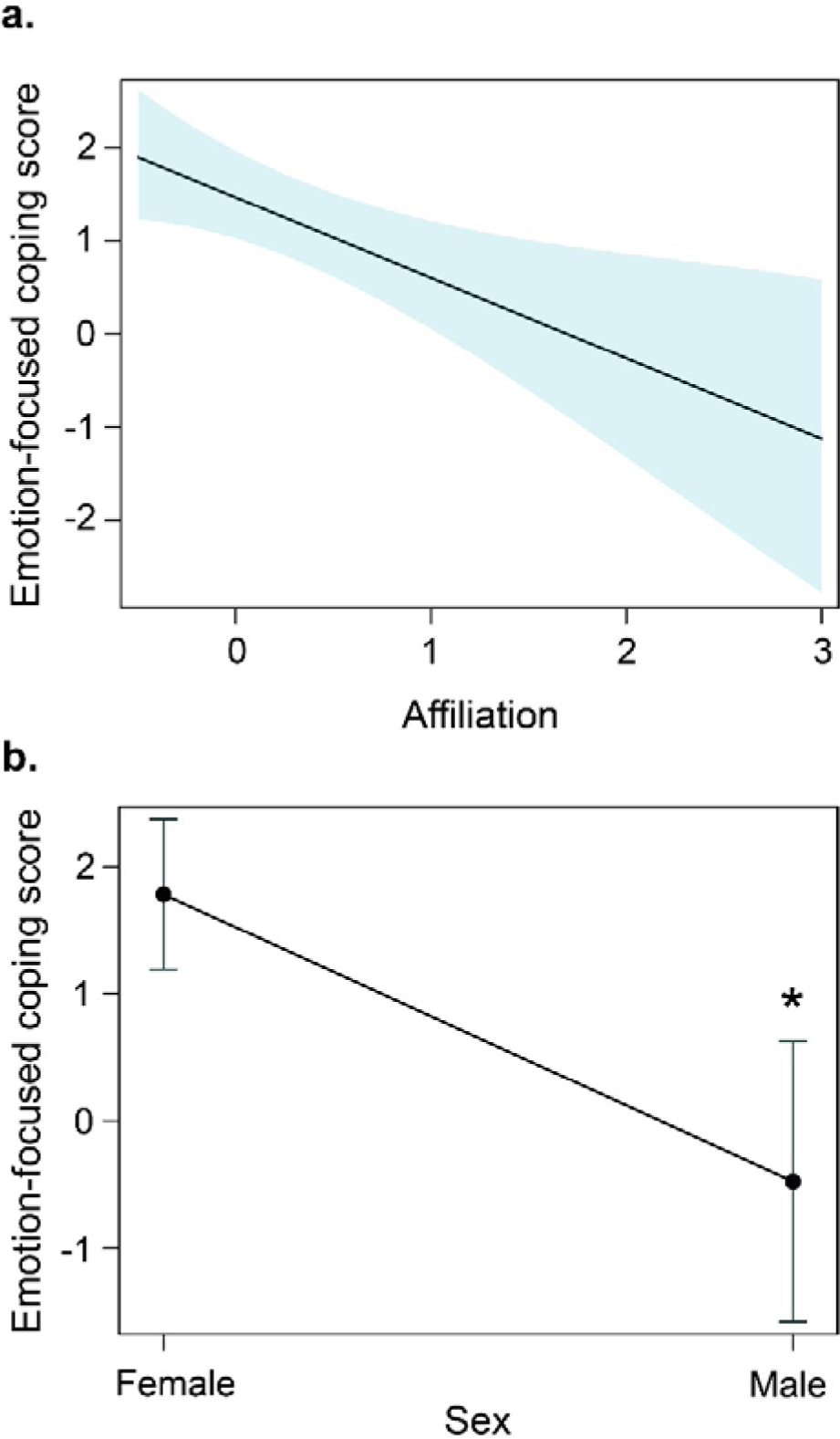
Effect of personality trait affiliation and sex on emotion-focused coping. **(a)** The effect plot shows an inverse relationship between emotion-focused coping scores and personality trait affiliation (LME: t = −6.00, *η^2^* = 0.95, 95% CI = 0.32, 1, p = 0.02, n=7). The shaded area in the figure shows the 95% confidence interval. **(b)** Females scored higher than males regarding emotion-focused coping (LME: t = −6.81, *η^2^* = 0.96, 95% CI = 0.42, 1, p = 0.02, n=7). The mean and the standard error values are shown in the figure.

No correlation was found between problem-focused coping and dominance rank (Spearman’s Rank Correlation: rho = 0.02, p = 0.95). We also found no correlation between emotion-focused coping and dominance rank (Spearman’s Rank Correlation: rho = −0.43, p = 0.42).

### COMT Val/Met polymorphism and behavioural coping styles

Sequencing results revealed two alleles equivalent to the human *COMT* Val^158^Met encoding polymorphism in long-tailed macaques. Out of the 26 individuals genotyped, 16 were homozygous A/A (Met encoding allele), six were heterozygous (A/G), and four were homozygous G/G (Val encoding allele). Accordingly, the targeted polymorphism was detected at 61.53%, 23% and 15.38% frequencies, respectively (allele frequencies: A = 0.73; G = 0.29).

We analysed a subset of individuals for whom *COMT* polymorphism information was available, and a specific coping style could be assigned (n = 18). Due to a small sample size, we could only compare homozygous A/A (n = 6) and G/G (n = 3) individuals from the problem-focused coping category. We found a significant difference between the coping scores of homozygous G/G and A/A individuals (Wilcoxon Rank-Sum test: z = −2.20, *η^2^* = 0.6, p = 0.02, **Fig. 4**). Homozygous G/G individuals (1.07 ± 0.19) scored significantly higher regarding problem-focused coping than their homozygous A/A counterparts (0.60 ± 0.15). This was in contrast to the personality traits, which did not correlate with the genetic polymorphism (see **Supplementary Fig. 4**). Finally, if we visually inspect the arousal data, it is interesting to see that the two individuals with the G/G genotype of which we also had thermal data were the ones that recovered most (marked by high nose temperatures) from the stressor (see **Fig. 2**).

**Fig. 4.**
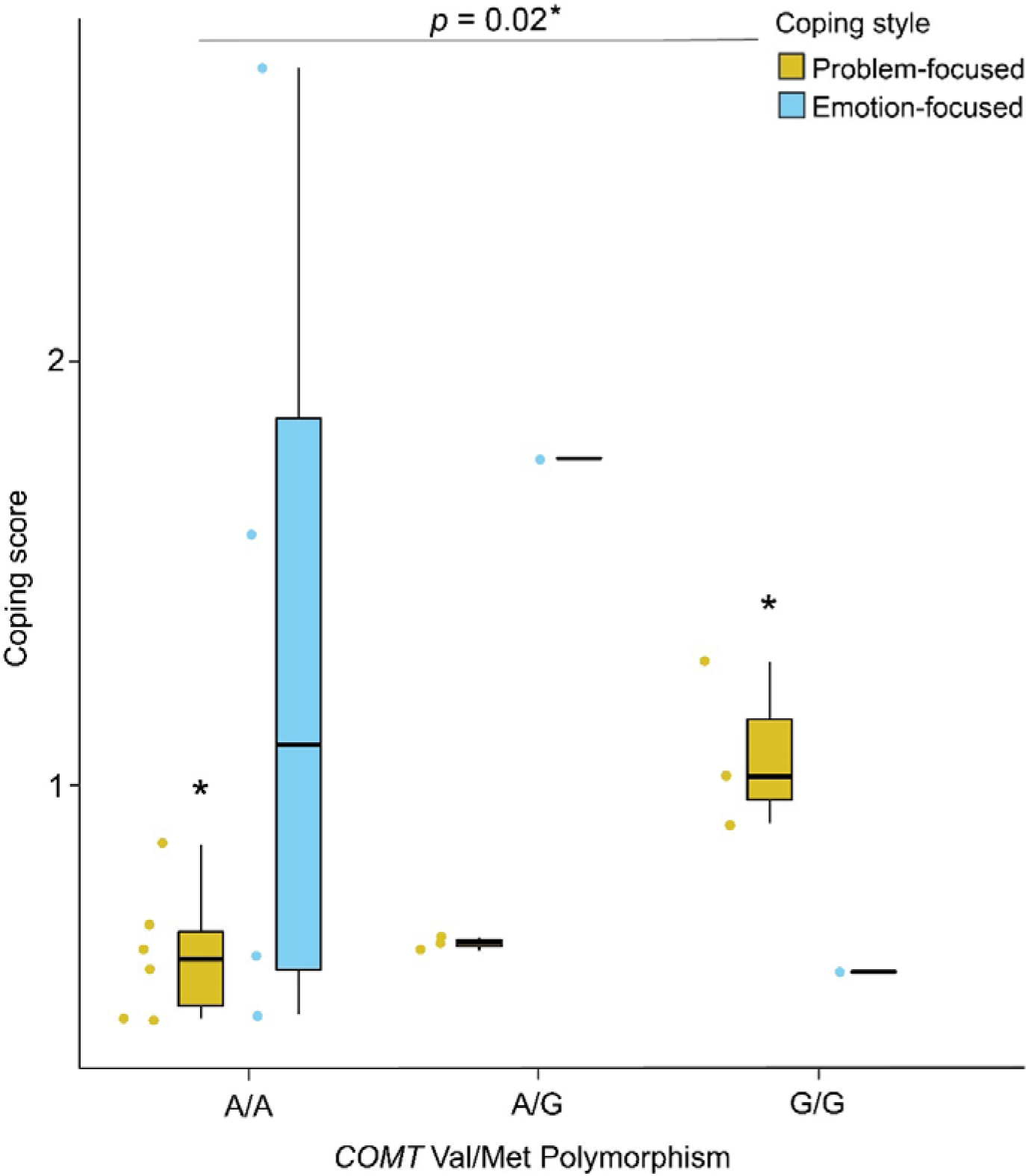
*COMT* Val/Met encoding polymorphism and behavioural coping styles. The box plot shows the two coping strategies and their association with *COMT* Val/Met genotype. A significant difference between the coping scores of homozygous G/G and A/A individuals was found (Wilcoxon Rank-Sum test: z = −2.20, *η^2^* = 0.6, p = 0.02, n=9). Individual data points are represented using solid dots. Boxes represent interquartile ranges, and whiskers represent the upper and lower limits of the data. The horizontal bars within the boxes represent the median values.

## Discussion

We investigated coping mechanisms in an NHP species using a multidisciplinary research design (See **Fig. 5** for an overview). Unlike a proactive-reactive continuum, non-correlating problem- and emotion-focused coping styles were identified, characterised by direct aggressive engagement with stressors and reliance on conspecifics for social support through affiliation, respectively. Problem-focused copers exhibited lower emotional arousal than emotion-focused ones, as examined using a non-invasive infrared thermal imaging method. Personality traits partially predicted coping, particularly the emotion-focused coping style, where higher affiliation was associated with lower emotion-focused coping scores. We furthermore provided the first evidence of the human equivalent *COMT* Val/Met polymorphism in long-tailed macaques. Although identified in a few individuals only, we could show that the G variant in a homozygous state (G/G, two alleles encoding for Val) was associated with problem-focused coping. This suggests a genetic basis for a complex behaviour. Our findings underscore the importance of multidisciplinary approaches in understanding coping mechanisms and their evolution.

**Fig. 5.**
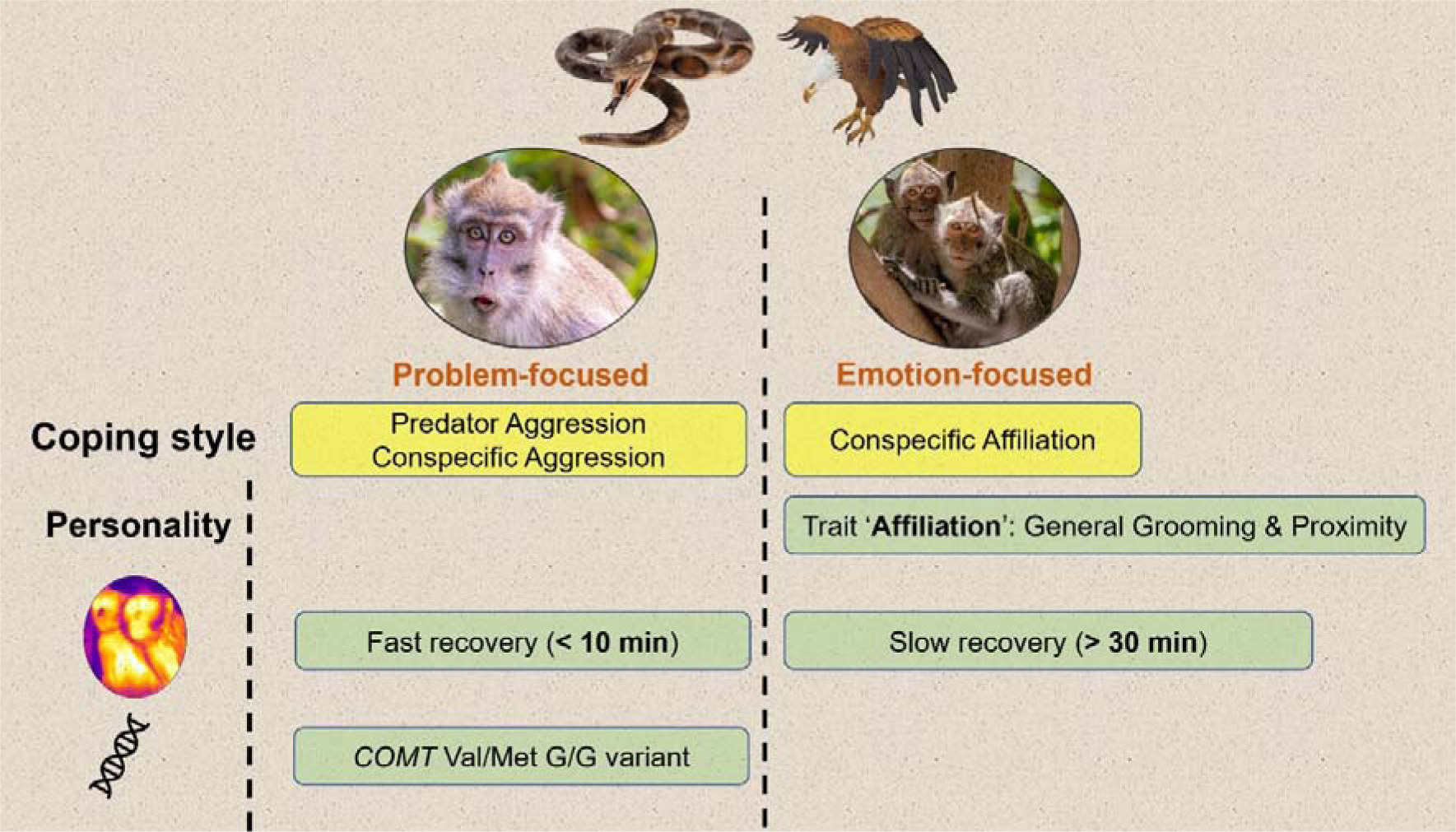
Overview of the study. Schematic figure showing the framework and findings of the current study.

Our results align with previous studies on coping styles and associated variations among individuals ^7,10,14,21,22,26^. However, unlike a proactive-reactive continuum ^10^, which is predominantly described in the non-human animal literature, we found non-correlating problem- and emotion-focused coping styles, suggesting independent coping strategies in long-tailed macaques (see ^52^). The problem-focused coping style was associated with direct engagement with the stressor through aggressive reactions and also aggression towards conspecifics. Out of all the individuals to whom a specific coping style could be assigned (n = 22), we found ∼68% of them to have problem-focused coping styles. This is in line with the idea that a despotic species, like long-tailed macaques, would exhibit lower inhibition control than socially tolerant species ^53^. Aggression towards conspecifics, on the other hand, can be attributed to ‘frustration’ and ‘redirection’, critical signs of societies with low social tolerance ^46,54^. The problem-focused copers were less emotionally aroused after an initial 10-min of predator exposure than their emotion-focused counterparts; it might indicate their predisposed neural underpinnings, which made them ‘proactive’ ^8^, i.e., better equipped to engage and cope with stressful situations. In other words, the problem-focused copers were more efficient in controlling (albeit not voluntarily) emotional arousal than emotion-focused copers with regard to the activities of the autonomic sympathetic nervous system. On the other hand, the emotion-focused coping dimension had positively loaded variables of close (distance to stressor) and conspecific affiliation and negatively loaded variable of far (distance to stressor). Even though the variables close and far were not mutually exclusive (see **Supplementary Table 1**), they seem to have covaried. Yet interestingly, it suggests that individuals with emotion-focused coping styles were not entirely avoiding the stressor but, at the same time, not directly engaging with it. These results suggest the complex and multifaceted structure of coping and the less distinctive extremes of its strategies ^9,23^. However, these individuals also seemed to rely on support seeking through affiliation from conspecifics, which might act as ‘social buffering’ ^55–57^. Thus, an emotion-focused coping strategy might also seem beneficial in long-tailed macaques, where close social bonds are prerequisites for maintaining group cohesion, cooperative interactions and sociality ^58,59^.

A relatively new and non-invasive measure of infrared thermography captured emotionality in coping in our study. As opposed to the problem-focused copers, heightened and persistent emotional arousal in emotion-focused copers was found. Thus the behavioural coping styles correlated with HPA axis reactivity. These findings corroborate that emotion-focused copers require prolonged social support from conspecifics to reduce arousal and cope with stressors. While the apparent non-independence of behavioural coping styles with HPA axis reactivity may seem contrasting to the prediction of the two-tier model, it should be noted that thermography data from the low reactants could not be obtained. Although they neither significantly engaged in predator aggression nor relied on conspecific support, the missing information on emotionality has made it challenging to conclude whether behavioural coping styles and reactivity are (in)dependent components of coping. Nevertheless, they seem to have covaried. In wild baboons, while personalities did not affect coping styles, neuroticism was predicted by stress reactivity ^60^. Finally, despite controlling for obvious confounding factors (e.g., indoor/outdoor temperature differences, distance and angle of the subjects from the thermal camera, etc.), other unprecedented factors (see ^61^) can cause a reduction in the precision of the thermal measures. Therefore, a methodological comparison between infrared thermography and endocrine measures would be recommended to better understand the HPA axis reactivity in coping.

Using the comprehensive multi-method approach of behavioural observations and experiments, we found three personality traits in long-tailed macaques – activity-sociability, affiliation, and exploration. Traits similar to activity-sociability and affiliation had previously been identified in long-tailed macaques ^62^. While activity-sociability and affiliation in our study represented variables from an observational perspective, repetitive ‘rare’ behaviours were captured using experiments in the exploration dimension. This validates the concept of objective assessment of personalities in non-human animals ^42,63^. Age and sex were found to influence personality traits. Age had a negative effect on activity-sociability and exploration. This effect may be attributed to the active and playful underpinnings of some loaded variables within those traits, e.g., *social play, handling container,* and *object manipulation*. Play behaviour is essential for the locomotive, cognitive, and social development of young individuals ^64^. Therefore, younger individuals are expected to score higher on these traits than older individuals, in which activity budgets and priorities may have shifted more towards goals such as reproduction and competition ^65^. Alongside age, sex was also found to significantly predict personality scores on activity-sociability and exploration, in which males scored higher than females. This may be attributed to female philopatry and male dispersal in long-tailed macaques, where males tend to leave their natal group around the age of four years and join either a bachelor or a new social group ^66^. Although personalities, following a strict definition, should not be sex-dependent, there is growing evidence for sex-specific personality dimensions. Differential selection pressures and varying life history trajectories are considered the underlying mechanisms for the evolution of sex-dependent personality traits ^67^.

We found partial support for our hypothesis of personalities predicting coping styles. The observed personality traits did not affect problem-focused coping styles. This indicates that the problem-focused coping style in itself may be an independent personality trait equivalent to boldness, and the associated variables of our problem-focused coping may have captured boldness-related behaviours. However, it is also important to emphasise that we did not find any evidence of ‘boldness-explorative’ behavioural syndromes, as reported in previous studies ^68,69^. Emotion-focused coping, on the other hand, was inversely associated with personality trait affiliation. This again supports the social buffering hypothesis that individuals with higher affiliative interactions are less prone to stress than individuals who scored low in the affiliation personality dimension. This furthermore strengthens the effectiveness of close social relationships in alleviating stress in group-living species. Interestingly, we noted that females scored higher than males in the emotion-focused coping dimension. While this observed sex difference may be explained by a general tendency of females to be more anxious than males ^70^, our results here should be interpreted with caution due to a low sample size of males. Contrary to our prediction, although linear and steep hierarchies in all three groups were noticed, we did not find any relationship between dominance-rank relationships and coping styles. A recent study on wild baboons (*Papio anubis*) also found no relationship between social hierarchies and coping styles ^71^. These results are unsurprising because dominance and coping styles are group- and individual-level properties, respectively, and might not always covary.

In our final analysis, we were interested in whether potentially different *COMT* genotypes modulate differences in the coping styles of long-tailed macaques. In humans, a functional polymorphism located in exon 4 of the *COMT* gene (Val^158^Met) has garnered significant scientific attention due to its counterbalancing effect on emotional resilience, stress, anxiety and cognition ^49^. Even though the catecholaminergic system appears highly conserved among vertebrates, the *COMT* gene has received little attention in NHP studies. To our knowledge, the Val/Met polymorphism has only been reported in Assamese macaques ^38^ among *Macaca* and thus has not been studied extensively regarding emotional arousal or coping styles in NHP species. We found an association between the *COMT* Val/Met polymorphism and problem-focused coping style; G/G individuals (Val/Val) scored higher on “offensive, direct, bold” coping than individuals carrying the A/A (Met/Met) variant, as hypothesised ^51^. This finding suggests a potential genetic basis for coping styles in long-tailed macaques ^72,73^. Our results align with the “warrior-worrier” model used to explain the existence of two alleles that induce distinct behavioural phenotypes in humans. The model claims that the Val-allele confers an advantage in confronting aversive stimuli (greater stress resilience and lower anxiety levels, i.e., warrior), whereas the Met-allele is beneficial in cognitive tasks (e.g., memory and attention tasks; worrier). Due to a low sample size, driven by just one individual carrying the G/G variant, comparisons could not be made among emotion-focused copers, which would be an important avenue for future studies. In addition, as highlighted, the two problem-focused copers with the G/G variant were least emotionally aroused, potentially indicating an effect of *COMT* Val/Met polymorphism on emotion. To what extent the *COMT* Val/Met polymorphism can influence animal emotion would thus be interesting to explore in future. It is, however, crucial to highlight that the same polymorphism had no relationship with the observed personality traits in this study, potentially indicating non-overlapping genetic underpinnings of coping styles (and potentially emotionality) and personalities.

The candidate genes of the serotonergic and dopaminergic systems are known to mediate coping in humans ^74^ and thus may explain the underlying causal mechanisms. However, as pointed out by earlier studies, there is an urgent need to consider the candidate genes while understanding the causal mechanisms of coping. Similar results in humans and long-tailed macaques regarding the *COMT* Val/Met polymorphism and coping further highlight an evolutionary basis of this mechanism and stress regulation in general. Although the effect of Val/Met on human behaviour has been intensively studied, we know that findings on the contribution of single candidate genes to a complex behavioural trait must be interpreted with caution ^75,76^. We call out for replication of our study, which is the first to reveal an impact of *COMT* on coping behaviour in an NHP species. Genome-wide association studies can help decipher the involvement of other gene variants, thereby providing a more detailed picture of the interaction of underlying genotype and coping behaviour. In addition, comparative studies on other species may reveal the evolutionary history of these traits and their genetic bases.

To conclude, we provided evidence of two different coping styles in a group-living NHP species using a comprehensive research design and identified the proximate mechanisms pertaining to behaviour, physiology (emotional arousal) and genetics. The complexity and dimensions of coping can be wide and should be investigated carefully. While varying concepts and methodologies are currently being applied, multidisciplinary research designs with novel methodologies are crucial to understanding the overarching process of coping. Although species-typical responses can be at play, identical research designs should be applied to a wide range of taxa to get a grip on the evolution of coping.

## Methods

### A. Subjects and Housing

We studied three groups of captive long-tailed macaques (*Macaca fascicularis*) housed at the Biomedical Primate Research Centre (BPRC) in Rijswijk, the Netherlands. The first group (Gr.1) consisted of 15 individuals; four were under the age of one year at the beginning of the study and were not included (n = 11). The remaining 11 individuals consisted of seven females and one male, all above the age of three years, and two females and one male between the ages of one and three years. The second group (Gr.2) originally consisted of 18 individuals but was reduced to 17 as a male was removed early during the study due to compatibility issues with other group members (n = 17). The remaining 17 individuals included ten females and one adult male above three years, and two females and four males aged between one and three years. The third group (Gr.3) comprised four males, all aged above three years (n = 4). A detailed description of the individuals is provided in **Supplementary Table 8**.

Gr.1 and Gr.2 had continuous access to indoor and outdoor enclosures with two or more interconnected compartments each. Gr.3, on the other hand, had a single indoor and outdoor compartment access. For Gr.1 and Gr.2, the indoor and outdoor enclosures were approximately 49 m^2^ and 183 m^2^, respectively. Gr.3 had an indoor and outdoor enclosure of about 3,55 m^2^ and 3,88 m^2^, respectively. Immediately adjacent to the enclosures were other groups of long-tailed macaques that were partly visible. However, the three study groups had no visual contact with each other. They were kept at distant buildings or in the same building but on two different floors (like Gr.1 and Gr.2). See **Supplementary Note 3** for additional details on the enclosure structures and diet of the animals.

### B. Ethical Note

The BPRC is accredited by the Association for Assessment and Accreditation of Laboratory Animal Care (AAALAC) and licensed to keep non-human primates for ethological and biomedical research. The institution follows high standards of animal welfare and refinement measures. Our study complied with all ethical regulations and guidelines for animal testing of BPRC’s Animal Experiments Committee and Animal Welfare Organisation (Animal Welfare Organisation/ IvD approval no. 019E). In addition, during the studies, at least one experimenter was present who was trained and certified (FELASA Accredited Laboratory Animal Science 066/19AF) to conduct animal experiments in accordance with the requirements of Article 9 of the Dutch Experiments on Animals Act.

### C. Data collection

#### Coping

We conducted predator model experiments repeatedly to investigate coping. Two lifesized predator models were used, and the experiments were conducted in two phases, with an interval of at least two months in between. The order in which the experiments were carried out across groups was randomised. The first model was a snake (160 cm long and resembling a python; **Supplementary Fig. 5a**), and the second was a bird of prey (resembling a hawk with a wingspan of 70 cm; **Supplementary Fig. 5b**); both of which are natural predators of long-tailed macaques in the wild ^77,78^. The snake model had a coiled posture, known to elicit threat responses more than a sinusoidal posture ^79^, whereas the bird had a standing, wings-up position. The models were placed out of reach of the individuals (∼60 cm away from the enclosures), in front of the indoor enclosures for Gr.1 and Gr.2, and in front of the outdoor enclosure for Gr.3. We made sure that the neighbouring groups had no visual access to the predator models, either by blocking the corridors with natural tree branches (Gr.1 and Gr.2) or placing the models inside a 180 cm x 70 cm x 70 cm wooden box with only the front exposed to the to be tested group (Gr.3). All groups were habituated to these modifications (without the predators) on the days before testing for at least 24 hours. Besides, on the days of testing, the immediate next enclosure compartments belonging to the other groups were emptied. This is standard practice during enclosure cleaning and maintenance at the BPRC; therefore, it is unlikely to cause any stress on the focal (and other) groups. The predator exposure experiments lasted half an hour and were recorded using two video cameras (Canon Legria HF G25 and Sony FDR AX100E 4K) mounted on tripods from different angles.

The facial (nose bridge, tip and mid-nose) temperature data of the subjects were collected using infrared thermography (cf. ^80^) throughout the predator exposure experiments. Facial temperatures were collected using two FLIR E96 (640 x 480 thermal resolution) thermal cameras by two experimenters who either sat on the ground (for Gr.1 and Gr.2) or stood (Gr.3) at least 1 metre away from the enclosures. The emissivity was set at 0.97; typically, a value of 0.97 or 0.98 is used for non-human primates ^81^. Reflective- and ambient temperatures and humidity measures were set at the thermal cameras before using them. We used two (for reliability) ThermoPro TP50 thermo-hygrometer devices to record ambient temperature and humidity. Any ambient temperature and humidity changes were noted and subsequently adjusted in the thermal cameras during the experiments. All subjects were habituated to the presence of the experimenters for over a month and at least a week to the thermal cameras. Three key criteria were followed to collect reliable facial temperature data to assess emotional arousal: (i) the distance between the experimenter and an individual subject is within 2 meters, (ii) an individual is facing the thermal camera at an angle not more than 50°, (iii) an individual is not moving abruptly and staying relatively still (cf. ^19,45,80,81^).

#### Personality

Personality was assessed using a multi-method bottom-up approach consisting of behavioural observations and novelty experiments. A multi-method process is expected to deliver a more complete view of personality objectively than each method separately ^82^. Following an extensive and standardised ethogram ^42^, we used a continuous focal sampling method for the behavioural observation part. To investigate the temporal consistency in behaviour, being a prerequisite for personality traits ^83^, focal observations were conducted in two different phases: for Gr.1, the first phase took place from March to July 2022 and the second from August to September 2022; in Gr.2, the first phase spanned between February and June 2022, and the second from August to November 2022. In Gr.3, focal observations were conducted over one relatively long temporal window from November 2021 to April 2022. These observations were later divided into two non-overlapping phases to check for temporal consistency. Each focal follow was 20 minutes long and recorded with a video camera (Canon Legria HF R806), handheld or mounted on a tripod. The order of sampling was pseudorandomised, and no individual was observed consecutively. Focal sampling took place 3-4 days a week, between morning (0900 and 1200 hours) and afternoon (1201 and 1500 hours) feeding schedules of the monkeys. Each focal animal was observed during a morning and an afternoon session on the same day, and both sessions were equally sampled.

The experimental approach entailed exposing the individuals in their social group setting to three categories of novelties - food puzzles, novel food items, and novel objects (**Supplementary Fig. 6**). Each category had two types (i.e., two types of food puzzles, two types of novel objects, etc.) to check for contextual consistency in behaviour. Similar to the focal observations, the personality experiments were conducted in two different phases, rendering the commencement of a total of 12 experiments per group. For Gr.1, the first phase of experiments occurred from January to February 2022 and the second from May to July 2022. For Gr.2, phase one was carried out between April and July of 2022 and phase two between August and September 2022. For Gr.3, the first phase occurred between December 2021 and February 2022 and the second from August to November 2022. Only one experiment was conducted on a given day; their order was pseudorandomised between the two phases. Experiments belonging to the same category were not conducted consecutively. In addition, observations and experiments were conducted on different days for a group. We performed all the experiments in the indoor enclosures except for Gr.3, where experiments took place in both indoor and outdoor enclosures. For each experiment, the animals were locked briefly in their outdoor (applicable to all) or indoor (applicable to Gr.3) enclosures while the task was being set up. At the BPRC, all macaques are trained to move to outdoor or indoor enclosures by voice commands of the caretakers. Therefore, movement before the experiments most likely caused no additional stress to the animals. All experiments were recorded using video cameras (Canon Legria HF R806, Canon Legria HF G25, and Sony FDR AX100E 4k) mounted on tripods. Each camera focused on a specific compartment of Gr.1 and Gr.2 enclosures. For Gr.3, due to its small enclosure area, two cameras were placed, one focusing on the indoor and one on the outdoor enclosure. Video recording was started before the animals regained full access to their enclosures and ended when the experiments were over.

We used two types of food puzzles: a box and a pipe. A wooden maze box (∼30 cm x 40 cm x 15 cm; **Supplementary Fig. 6a**) with a plexiglass front cover was used, which had two narrow horizontal openings, enabling individuals to move the food rewards (cut pieces of apples) with their fingers. Solving the puzzle required animals to move the reward far enough to the left so that it would roll downwards to the right. At that point, it would have to be moved to the left again towards the opening from where it could be retrieved. During each experiment, three identical boxes containing multiple cut pieces of apples were secured to the bars of the indoor enclosure, roughly 15-20 cm above the ground. The distance from the ground allowed enough space for the apples to fall out. For Gr.3, considering the smaller group size, we used two boxes instead of three. Multiple box puzzles (similarly applicable to other objects/items) helped reduce monopolisation.

The pipe puzzle (∼70 cm in length, Ø 15 cm; **Supplementary Fig. 6b**) was made of hard plastic and had a row of small holes (∼Ø 2 cm) on the upper side. During each experiment, three and two identical pipes, for Gr.1/Gr.2 and Gr.3, respectively, containing three handfuls of a seed mix (corn, sunflower, etc.), were hung up roughly 100 cm from the ground. The food rewards could be retrieved by rotating and holding the pipe far enough for the rewards to fall downwards through the holes. The tension of the strings assisted the pipes to return to their original positions when released. Apart from multiple identical items, the puzzles were placed in a way (e.g., in different compartments) that potentially minimised monopolisation by higher-ranking individuals. The experiments started with monkeys regaining access to the enclosures and ended either after one hour or after all puzzles had been solved (i.e., all rewards were obtained), whichever was earlier.

The novel food items used during the first phase of Gr.1 and Gr.3 were rambutans (**Supplementary Fig. 6c**) and dragonfruits (**Supplementary Fig. 6d**). However, due to veterinary advice and reconsideration, the rambutans were replaced with starfruits during the second phase of experiments. The monkeys had no experience with either of these fruits. In Gr.2, starfruit and dragonfruits were used in both phases. During each experiment, multiple pieces of either intact (rambutans) or halved (dragonfruit and starfruit) fruits were placed in the enclosure ground. The number of novel food items was adjusted according to the group sizes to avoid monopolisation. Like food puzzles, the experiments started with monkeys regaining access to their enclosures and ended after one hour or when all items were eaten.

The two novel objects used were blue and green coloured plastic egg containers (14 cm x 13 cm x 4 cm; **Supplementary Fig. 6e**) and wooden massage rollers (24 cm x 10 cm x 4 cm; **Supplementary Fig. 6f**). For Gr.1 and Gr.2, six identical objects were used during each experiment and spread about 100 cm apart from each other in the indoor enclosure compartments. In comparison, we used two objects of each type for Gr.3. Since the individuals could carry the objects freely, we recorded the monkeys’ activities in indoor and outdoor enclosures. Each experiment lasted for an hour.

#### Collection and storage of blood samples for genetic analyses

EDTA whole blood samples were obtained from the monkeys during annual veterinary health check-ups to isolate genomic DNA (gDNA) using a standard salting-out purification procedure. The check-ups are distributed over the year and have the target of sampling each animal with an interval of approximately 12 months. The individuals in our study did not show any clinical signs of diseases based on daily care and observations. Besides, monkeys were only included in the annual veterinary health check-ups when they were ≥ 9 months old. The collection of blood samples took place in the morning, and monkeys did not receive food after 1700 hours on the previous day. Progressive refinement procedures were followed to minimise the stress of capture before the sedation process. Monkeys were trained to voluntarily enter the squeeze cage as a part of the refinement procedure. After an intramuscular injection of the sedative, blood sampling was carried out by qualified animal caretakers within a window of 20 minutes. The collection site was sterilised using 70% alcohol. The monkeys gained consciousness within 2 hours of the procedure. Notably, no behavioural experiments or observations were conducted within a week of the health check-ups.

From each monkey in the studied cohort, 30 µl of gDNA with a concentration varying between 46-649 ηg/μl (on average 124,5 ηg/μl) was shipped to Affenberg Landskron Field Research Lab (field research station of the University of Vienna; Landskron, Carinthia, Austria) for further analysis.

### D. Data analyses and statistics

#### Behavioural Coping styles

To investigate whether the predator models indeed played the role of stressors, we first compared self-directed behaviours (autogroom and scratch combined) between predator exposure experiments (all phases and predator models combined) and regular focal data (phases combined). Frequencies of self-directed behaviours were calculated per minute at the individual level. We calculated the frequencies from focal data when no aggressive interactions were observed for at least five minutes to eliminate any potential effect of social stress.

Videos were coded using Solomon Coder (version beta 17.03.22). The durational (s/hour) and event behaviours (frequencies/hour) were calculated per individual and were corrected for the time spent out of sight. We set up a minimum threshold of 6 minutes of within-sight observation (10% of the total combined observation time/phase) for the individuals to be considered for analyses. Despite attempts to record all individuals with multiple cameras, we could not obtain adequate data from two individuals (two females above three years of age) belonging to Gr.2. Therefore, we revised our sample size for the behavioural part of coping from 32 to 30 individuals. In addition, even if two predators could elicit responses from monkeys quite differently (see ^43^), we decided to combine their data for each phase (at the individual level) to tackle the low occurrences of some of the variables. Three experimenters coded the behavioural coping videos, and inter-rater reliability was calculated was found to be high ((ICC (3,k) = 0.97, p<0.001).

The consistency of the behavioural coping-related variables was examined using a two-way mixed model intraclass correlation (ICC (3,1)) analysis. Only variables with sufficient temporal consistency (ICC values ≥ 0.3 and p < 0.05) were included in further analyses. ICC analysis yielded 9 repeatable variables with ICC values ranging from 0.32 to 0.80 (**Supplementary Table 9**). These repeated variables (average values between phases one and two) were used in the EFA with a principal axis factoring (PAF) extraction method to test for the presence of coping styles. PAF was chosen as it does not assume a multivariate normal distribution, and the data might exhibit significant deviation from a normal distribution. After an initial run of the EFA, four variables, *foraging, locomotion, freeze,* and *yawn,* were removed due to low Kaiser-Meyer-Olkin measure of sampling adequacy (MSA) scores. The final model included 5 variables with an overall acceptable MSA score of 0.62. Bartlett‘s score of sphericity was significant (x^2^ = 41.56, p<0.001); thus, the assumptions of EFA were met. The latent factors were then extracted based on eigenvalues >1. The root mean square of the residuals showed a value of 0.02, which indicated a good model fit. In addition, a Tucker-Lewis Index (TLI) value of 0.95 was obtained, further suggesting that the model fit was sufficient ^84^. We performed a promax rotation method for meaningful interpretation of the potentially correlated factors and factor loadings. The resulting factors were labelled based on which behavioural variables loaded significantly (≥ 0.5, positive and negative) onto them and whether their nature was related to problem- or emotion-focused responses (i.e., coping styles ^10,23,26^). The scores of each individual per factor were extracted, and individuals were assigned to specific coping styles accordingly.

#### Emotional arousal

Following the three criteria of collecting thermal images, we extracted information from the individuals across all phases and predator models. The number of thermal images and individuals was adjusted based on the specific coping styles of the monkeys. One experimenter coded all the thermal recordings; another experimenter coded 5% of the videos to check for reliability; it was found to be high ((ICC (3,k) = 0.93, p<0.001). Not all individuals were equally sampled, as the procedure heavily depended on monkeys approaching the enclosure fences. The valid thermal images were imported to FLIR Tools (version 6.4.18039.1003) for analysis. Nose-bridge, tip, and mid-nose temperatures were extracted from each image. Each image was magnified (606%) consistently, and an area size of 2 x 2 pixels was applied to the designated regions to extract temperature measures (**Fig. 6**). Temperature measures from the three regions were then checked for correlation using Spearman Rank correlation tests. High correlations between the nose bridge, tip, and mid-nose regions for facial temperature readings (Spearmańs Rank Correlation: nose tip and mid-nose, rho = 0.99, p < 0.001; nose-tip and nose bridge, rho = 0.99, p < 0.001, mid-nose and nose bridge, rho = 0.99, p < 0.001) was found. Since the mid-nose region is less susceptible to temperature changes due to breathing, mid-nose temperature data were used for the analysis (see ^80^).

**Fig. 6.**
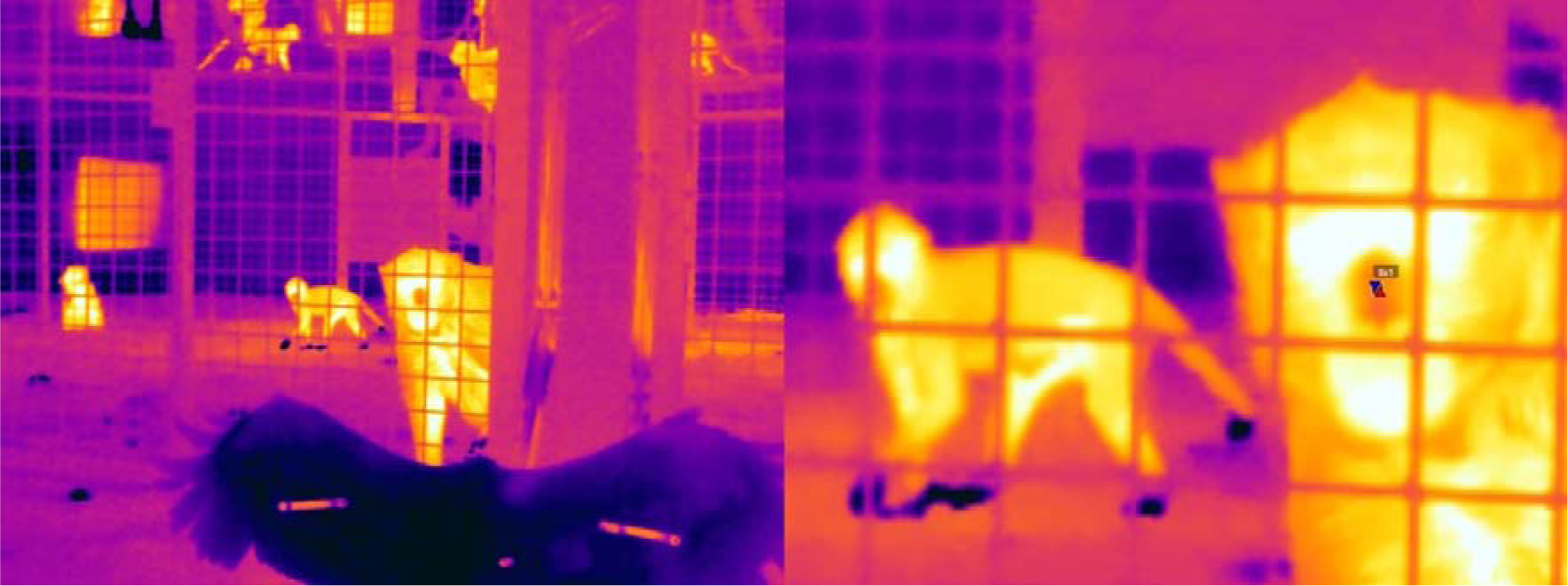
Infrared thermal imaging to measure emotional arousal. The left panel shows the experimental set-up using the predator hawk in one of the long-tailed macaque groups. The right panel shows a zoomed-in version (606%) where the mid-nose temperature is extracted from a focal individual using a 2 x 2-pixel area. The pixel area provides an upper (up arrow), a lower temperature (down arrow), and an average estimate. We used the average value for analysis.

Even though the animals were trained to move between in/out-door enclosures following the commands of caretakers, we could not discard the possibility of emotional arousal associated with this process. Therefore, instead of using potentially unreliable thermal measures (as control) before the introduction of the predator, we decided to divide the 30-min predator exposure period into small time windows for comparison. We did not code data for the first two minutes of the predator exposure to avoid any potential influence of temperature differences between the enclosures and emotional arousal due to the associated lock and release. Additionally, if an individual travelled, during the experiment, between the indoor and outdoor enclosures, a 1-min no coding window for the individual was applied to control for potential temperature differences; besides, data were not coded for 1-min if a focal individual was found vocalising (cf. ^19,45,80^). We divided the overall duration of the experiments into the following time windows: 2-10 min, 10-20 min, and 20-30 min. Such a division ensured maximised sampling effort from the individuals. Individuals were only included in the analysis if at least one valid data point was collected from each time window. In case multiple data points were found pertaining to one individual within a time window, the average value was used. Since the aim was not to determine how quickly the facial temperature changed upon predator exposure but rather how coping strategies worked throughout the session, we converted all temperature values of the individuals to ‘0‘ from the 2-10 min window. This was used as the baseline measure. However, this is not to say that we did not expect a potential drop in temperature upon predator exposure. Temperature changes in the following periods (i.e., 10-20 min and 20-30 min) were compared with the baseline. Finally, we conducted a validation test using locomotion and foraging behaviours to control for the potential confound of the general activity of the monkeys. Following the inclusion criteria, we could extract reliable data on emotional arousal from 12 individuals (six each for problem- and emotion-focused coping styles).

#### Personality

We collected data on all 32 individuals. A total of 10,229 minutes of focal data (Phase 1: mean ± standard deviation = 159.76 ± 2.67 observation minutes/individual; Phase 2 = 159.89 ± 2.64 min/individual) was collected in addition to the 36 personality experiments from the two phases. Four coders coded observational and experimental videos, and inter-rater reliability was estimated using a two-way mixed ICC coefficient (ICC 3,k = 0.96, p<0.001). We used 5% of the observational and experimental data to calculate the inter-rater reliability.

For each phase of the focal observation, durational variables (sec/min) and event behaviours (frequencies/min) were calculated per individual, and the varying observation minutes were corrected. The variables for each category of the personality experiments were coded (also see ^42^) as follows: *latency to approach* (sec) as the time it took for an individual to move from an initial 5-meter distance to a 1-meter radius of the novel object, novel food, or food puzzle. If no approach was made, *latency* was scored as the total length of the experiment; *proximity* (s/hour) as the total time an individual spent in proximity (≤ 1 meter) of the novel object, novel food, or food puzzle; *manipulating* (s/hour), or *handling* (s/hour), as the time an individual spent manipulating or handling the food puzzle or novel object, respectively. The coding of *proximity* was seized as soon as an individual began *manipulating* or *handling* the experimental items. All variables were standardised for meaningful interpretation of the next steps of analyses.

All three groups were used together to view the species-level personality constructs comprehensively. The temporal consistency of the variables was tested using a two-way mixed model ICC, which compared each observational and experimental variable between phases one and two. A conservative cutoff was set in which only variables with ICC values ≥ 0.5 and p < 0.05 were considered temporally repeatable. Variables in which over half (i.e., more than 16 individuals) of the individuals had zero occurrences were excluded.

We found 33 repeatable variables from the ICC analysis with values ranging from 0.50 to 0.95. The average values of these variables between phases one and two were included in a PCA, but the number was further decreased to 17 after removing those with low communality scores (cut-off value = 0.7) and to even 12 after removing those loading in more than one component with similar magnitude. The communality scores of the remaining 12 variables ranged between 72.28% and 94.14%, with an overall Kaiser-Meyer-Olkin measure of sampling adequacy (MSA) value of 0.81 (see **Table 1**). A scree plot was generated using an unrotated PCA, and the eigenvalue of each potential principal component was reviewed alongside the percentage of variance explained individually and cumulatively. The number of principal components was subsequently decided on by inspecting the plot, eigenvalue scores (>1), and the amount of variance explained cumulatively (>70%). Since personality traits can theoretically be correlated and form behavioural syndromes, we used a direct oblimin rotation technique. Factor loadings ≥ 0.5 (for both positive and negative loading) for the variables were considered significant. Finally, variables were removed if loaded on multiple components with similar magnitude. The resulting principal components were labelled based on the significant behavioural variables loaded on them. The scores of each individual per component, or personality trait, were extracted. Effects of sex and age on the personality traits were investigated.

#### Dominance hierarchy

Dominance rank relationships were calculated at the group level using a Bayesian Elo-rating method ^85^. The method was based on calculating winning probabilities, and we particularly used submissive behaviours coded from focal observations (*avoid, be displaced, fear grimace, flee,* and *social presence*, see ^42^) to assess the steepness of hierarchies. We checked for the independence of the above behaviours from personality assessment. After constructing the hierarchies, individuals were plotted according to their respective estimated Elo values (**Supplementary Fig. 7**).

#### COMT Genotyping

We genotyped our study animals (*n=26*) for the equivalent site of the human Val^158^Met encoded polymorphism located in exon 4 of the *COMT* gene. However, in macaques, the *COMT* gene encodes a 270 amino acid (aa) long product, which is one aa shorter as compared to the human COMT protein. As such, the human *COMT* Val^158^Met encoded polymorphism in macaques is located at codon 157 ^41^. Amplification of the target site and sample preparation for sequencing was conducted at the Affenberg Landskron Field Research Lab (field research station of the University of Vienna; Landskron, Carinthia, Austria). Amplification products were outsourced for sequencing to LGC genomics (Berlin, Germany).

We used primers of Pflüger et al. (^40^) to amplify a 288 base pair product covering the target site (forw: 5′-AAGATCGTGGACGCCGTG, rev: 5′-ACAGTGGGTTTTCAATGAACGTG). Amplification was performed in duplicates along with two non-template controls using a MIC qPCR cycler (Bio Molecular Systems, Coomera, Australia). The 20 µl qPCR reaction contained nuclease-free H_2_0 (ThermoFisher Scientific, Geel, Belgium), Buffer B2 (1x conc., Solis BioDyne, Tartu, Estonia), 2.5 mM MgCl_2_ (Solis BioDyne, Tartu, Estonia), 0.2 mM dNTP’s (Solis BioDyne, Tartu, Estonia), 0.2 µM of each primer (Integrated DNA Technologies, Leuven, Belgium), EvaGreen (1x conc, Biotium, Inc. 46117 Landing Parkway Fremont, CA 94538), 0.05 U/µl hot-start DNA polymerase (HOT FIREPol® DNA polymerase, Solis BioDyne, Tartu, Estonia), and 2 μl template DNA per reaction tube. Reactions were prepared in a sterile UVP PCR2 Workstation (Analytik Jena GmbH, Jena, Germany).

We cycled 15 min at 95 °C, following 40x 10 sec at 95°C, 40 sec at 60 °C and 20 sec at 72 °C. To ensure that only a single product resulted from amplification, we included a melting curve analysis ranging from 72°C to 95°C with 0.3 °C/s. The experimental samples showing a single peak at the expected melting temperature were purified using the Omega E.Z.N.A. Gel extraction kit (Omega Bio-tek, Inc. Norcross, USA). The spin column was loaded with 20µl PCR sample mixed with 20µl binding buffer. Duplicates of the same experimental sample were pooled. For purification, we followed the manufacturer’s instructions with only slight modifications. We added another washing step and refrained from loading the samples on an agarose gel to secure DNA yield. We used the melting curve results to verify the amplification products, and 10 µl of each sample plus 4 µl of forward primer were sent to LGC genomics. The resulting Sanger sequencing electropherograms were evaluated using the CodonCode Aligner software (version 10.0.2; CodonCode Corporation, Dedham, USA). Three additional non-synonymous SNPs were identified downstream of the target site. These polymorphisms were not linked to the target and, therefore, not subjected to further analyses.

#### Statistical models and packages

All statistical analyses were conducted in R (version: 4.3.1) ^86^. Generalised linear mixed-effect (GLMM) and linear mixed-effect model (LME) analyses were conducted using the *lmerTest* and *glmmTMB* packages. Null versus full model comparisons were done for all the models using the ‘lrtest’ function of the package *lmtest*. Model diagnostics were checked using *DHARMa* package. The multicollinearity of the fixed effects was examined with the help of the *performance* package, and a variance inflation factor of < 4 was set as a threshold for low correlation. Model selection was made based on the Akaike Information Criterion (ΔAIC = 2 as threshold) value using the ‘anova’ function. The significance value was set as 0.05 for all statistical tests, except for post hoc pairwise comparisons, where adjusted p-values were used following a Bonferroni correction method. See **Supplementary materials** for details on statistical model outputs.

## Data availability

Data will be made available upon publication.

## Acknowledgements

We thank Sophie Waasdorp, Charlotte Kluiver, and Sjoerd Sijbrandij for their assistance during the experiments and organisation of the data. We are grateful to Elisabeth H.M. Sterck and Jan A.M. Langermans for their help in implementing the study at the BPRC. We thank all the caretakers at the BPRC for their help and assistance. We acknowledge the Affenberg Zoobetriebsgesellschaft mbH for providing student housing and bearing operating expenses during lab work. We would also like to thank Martin Hofer and Ron van Sambeek for their support during lab work at the Affenberg field research lab and infrared thermal cameras, respectively. The study received funding from the European Union’s Horizon 2020 Marie Skłodowska-Curie Actions research and innovation program under grant number H2020-MSCA-IF-2019-893016 (awarded to D.B.).

## Contributions

D.B., A.R.G., and J.J.M.M. conceived and designed the study; D.B., A.R.G, P.E.C., and E.M. performed research, collected, coded and organised data; D.B., P.E.C., J.J.M.M., and L.S.P., managed and coordinated the study; N.G.G. provided the DNA samples; J.J., B.W., and L.S.P. carried out the genotyping work; D.B. and A.R.G. conducted the formal analysis of the data; D.B. wrote the first draft of the manuscript; D.B., N.G.G., J.J.M.M, and L.S.P. edited the manuscript. J.J.M.M. and L.S.P. supervised the study.

