## Supplementary Material for "Behavioural, physiological, and genetic drivers of coping"

**Table of contents**

| Item | Description | Page |
| --- | --- | --- |
| Supplementary Fig. 1 | Frequency of self-directed behaviour | 02 |
| Supplementary Table 1 | Ethogram of coping-related behaviours | 03 |
| Supplementary Fig. 2 | Categorisation of individuals into coping styles | 04 |
| Supplementary Note 1 | Group-level details of coping styles | 04 |
| Supplementary Table 2 | Effect of coping styles on nose-temperature change | 04 |
| Supplementary Fig. 3 | Average mid-nose (absolute) temperatures during predator exposure experiment | 05 |
| Supplementary Note 2 | Effects of age and sex on personality traits | 05 |
| Supplementary Table 3 | Effect of age and sex on activity-sociability | 05 |
| Supplementary Table 4 | Effect of age and sex on exploration | 06 |
| Supplementary Table 5 | Personality scores of the individuals | 07 |
| Supplementary Table 6 | Effect of personality on problem-focused coping | 08 |
| Supplementary Table 7 | Effect of personality on emotion-focused coping | 08 |
| Supplementary Fig. 4 | *COMT* Val/Met polymorphism and personality | 09 |
| Supplementary Table 8 | Details of the monkeys in the study. | 10 |
| Supplementary Note 3 | Additional details on enclosure and diet of the animals | 11 |
| Supplementary Fig. 5 | Predator models used in the study | 11 |
| Supplementary Fig. 6 | Novelty experiments for personality assessment | 12 |
| Supplementary Table 9 | ICC results on coping-related variables | 13 |
| Supplementary Fig. 7 | Dominance hierarchies of the groups | 14 |

**Supplementary Fig. 1**

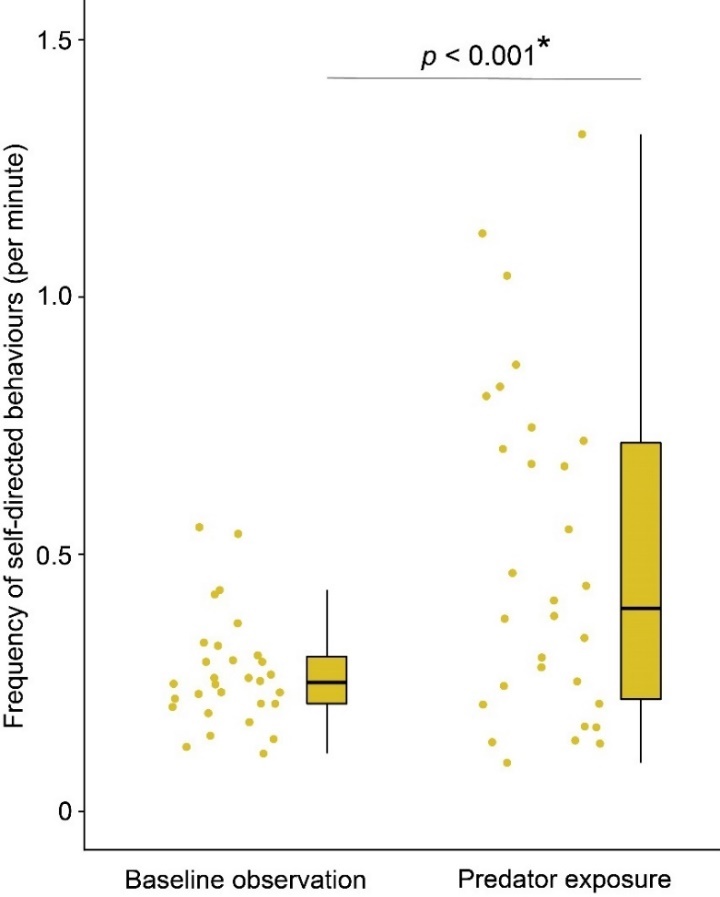

**Supplementary Fig. 1 – Frequency of self-directed behaviour.** The box plot shows the frequency of self-directed behaviour (per minute) during baseline observation and predator exposure (Wilcoxon Signed-Rank test: z = -3.42, r = 0.66, p < 0.001, n=30). Individual data points are represented using solid dots. Boxes represent interquartile ranges, and whiskers represent the upper and lower limits of the data. The horizontal bars within the boxes represent the median values.

**Supplementary Table 1**

| **Behaviour** | **Type** | **Definition** |
| --- | --- | --- |
| **Enclosure use** | | |
| Close ground | State | Individual is within a 1-meter radius of the stressor and on the ground. |
| Far | State | Individual is outside of the 1-meter radius from the stressor and located either on the ground or on hanging structures of the enclosure. |
| **Activity** | | |
| Locomotion | State | Individual walks or runs around, moves from one location to another, and is not stationary for more than 3 seconds. |
| Foraging | State | Individual moves slowly while looking for food on the ground, or sits/stands while looking for food on the ground. Also includes active eating/ingesting/handling of food items. |
| **Aggression** | | |
| Conspecific aggression | Event | Open-mouth threat: Individual opens his mouth for a while, directed at the receiver of aggression. Chin often pointed forwards.  Chase: Individual runs after a conspecific for at least three seconds.  Lunge: Individual jumps a maximum of two body lengths towards a conspecific.  Stare: Individual looks intensely at a conspecific with a straight back and raised eyebrows in an attempt to threaten.  (any one of the above qualifies for conspecific aggression) |
| Predator aggression | Event | Stare: Individual looks intensely at the stressor with straight back and raised eyebrows for at least 3 seconds.  Vocalisation: Individual makes loud warning calls and barks at the stressor. The direction of the head always remains toward the stressor. During the process, the erection of body hair or piloerection can be seen.  (any one of the above qualifies for predator aggression) |
| **Self-directed behaviours** | | |
| Autogroom | Event | Individual grooms oneself or closely inspects skin, nails, hands, toes or other body parts. |
| Scratch | Event | Individual uses fingers, hands, or foot to rake across own skin. |
| Freeze | Event | Individual maintains a tense body posture with no movement or vocalisations for at least 3 seconds. |
| Yawn | Event | Individual opens mouth wide and inhales intensely, which can be seen by the expansion of the chest. |
| **Conspecific affiliation** | | |
| Groom and lip-smack towards conspecifics | State and  Event | Groom: Individual touches and strokes another individual’s fur gently with one or both hands, accompanied by periodic hand contact with their own mouth. Individual pays close attention to the recipient’s fur (state).  Lip-smack: Individual opens and closes their lips rapidly, accompanied by a smacking sound, directed at a conspecific (event).  (any one of the above qualifies for conspecific affiliation) |

**Supplementary Table 1 – Ethogram of coping-related behaviours**. Name, type, and definition of all coping-related behaviours.

**Supplementary Fig. 2**

|  | Reactive |
| --- | --- |
| -1.76295 | -1.26708 |
| -0.142055 | -0.292442 |
| -1.12908 | -2.26369 |
| -1.71648 | -0.521106 |
| -0.477371 | -0.228951 |
| -0.477202 | -0.1017 |
| -0.455075 | -0.65637 |
| 0.634248 | 0.385975 |
| 0.635208 | 0.163783 |
| 0.6719 | 0.388807 |
| 0.572192 | 0.0224816 |
| 1.29235 | 0.115393 |
| 1.02027 | 0.720073 |
| 0.446211 | -0.707406 |
| 0.861143 | -0.287061 |
| 0.612007 | -0.585843 |
| 0.909914 | -0.856997 |
| 0.61499 | -0.979817 |
| 0.066587 | -0.189153 |
| 0.224152 | -0.220327 |
| 0.652382 | -0.0595051 |
| 0.304916 | -0.446293 |
| 0.446213 | -0.124191 |
| -0.531291 | 1.77044 |
| -1.8201 | 0.303627 |
| -0.251463 | 0.600597 |
| -0.394157 | 1.59396 |
| -0.371427 | 0.46419 |
| -0.416472 | 2.6941 |
| -0.0195606 | 0.564503 |

**Supplementary Fig. 2 – Categorisation of individuals into different coping styles.** Individuals (n=30) were plotted and categorised into problem-focused-, emotion-focused- and mixed coping styles based on scores obtained from the exploratory factor analysis. Individuals with low reactivities were also seen and represented. One individual (highlighted using a solid red dot), even though grouped into problem-focused, did not score higher than the population median (scores: problem-focused factor = 0.07, population median = 0.15). Subsequently, the individual was grouped as a low-reactant for analysis.

**Supplementary Note 1 – Group-level details of coping styles.**

At the group level, we found variations in percentages of problem- and emotion-focused copers. In Gr.1, ~36% of the individuals had problem-focused coping styles, ~53% had emotion-focused coping styles, and ~0.9% had low reactivity. Gr. 2 had ~53% problem-focused copers and, interestingly, no emotion-focused copers, but ~46% of the animals had low reactivity. We found 75% and 25% of the monkeys in Gr.3 to have problem-focused- and emotion-focused coping styles, respectively.

**Supplementary Table 2**

*Model:* Temperature change ~ coping style + time window + sex + (1|Group/id)

| **Fixed effects** | **Estimate** | **Std. error** | **df** | **t value** | **p-value** |
| --- | --- | --- | --- | --- | --- |
| (Intercept) | -0.204 | 1.464 | 4.114 | -0.140 | 0.895 |
| Time window (10-20 min) | 1.264 | 0.696 | 22.00 | 1.815 | 0.083 |
| Time window (20-30 min) | 1.315 | 0.696 | 22.00 | 1.889 | 0.072 |
| Coping style (emotion-focused) | -2.291 | 0.920 | 7.676 | -2.490 | 0.038* |
| Sex | 1.565 | 1.112 | 8.349 | 1.408 | 0.195 |

*(Significance code:* 0.01 ‘*’*)*

**Supplementary Table 2 – Effect of coping styles on nose temperature change.** Summary of the linear mixed-effect model.

**Supplementary Fig. 3**

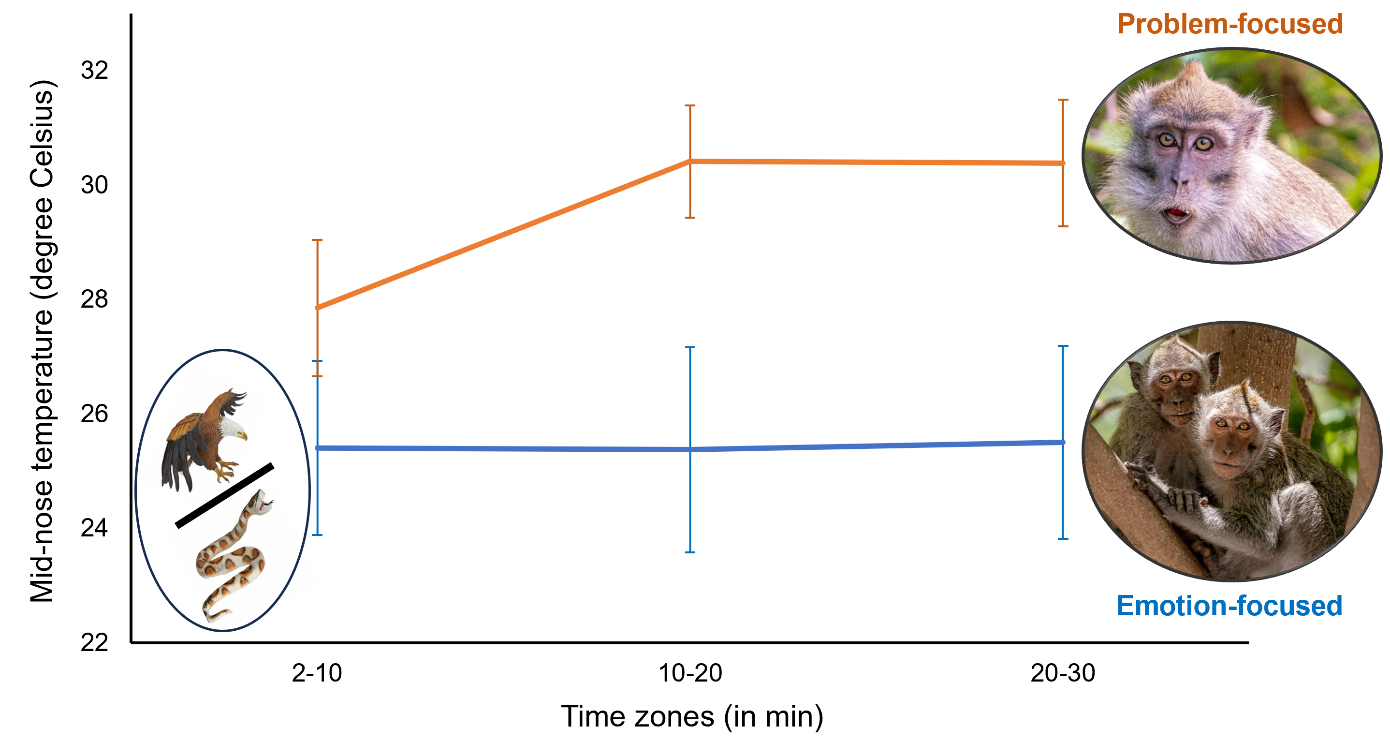

**Supplementary Fig. 3 – Mid-nose temperatures of the problem- and emotion-focused copers at different time windows of the predator exposure experiments.** The absolute average (± standard error) mid-nose temperatures of the problem- and emotion-focused copers (n=12) during predator exposure experiments.

**Supplementary Note 2 – Effects of age and sex on personality traits.**

We did not find any effect of age and sex on affiliation; however, independent effects of age and sex were found on activity-sociability and exploration. We found a significant negative effect of age on activity-sociability (GLMM: p = 0.006, **Supplementary Table 3**) and exploration (GLMM: p < 0.001, **Supplementary Table 4**). Males were, furthermore, found to score higher than females in both activity-sociability (female: -0.42 ± 0.66; male: 0.70 ± 1.10; GLMM: p = 0.002, **Supplementary Table 3**) and exploration (female: -0.37 ± 0.73; male: 0.62 ± 1.10; GLMM: p = 0.02, **Supplementary Table 4**) personality traits.

**Supplementary Table 3**

*Model:* Activity-Sociability ~ Age + Sex + (1|Group/id)

| **Fixed effects** | **Estimate** | **Std. error** | **z value** | **p-value** |
| --- | --- | --- | --- | --- |
| (Intercept) | 0.178 | 0.274 | 0.648 | 0.516 |
| Age | -0.096 | 0.035 | -2.722 | 0.006 ** |
| Sex (Male) | 0.865 | 0.285 | 3.027 | 0.002 ** |

*Comparison with null model:* Likelihood ratio test - χ2 = 18.17, p < 0.001 ***

*(Significance code:* 0.01 ‘*’, 0.001 ‘**’, 0.000 ‘***’*)*

**Supplementary Table 3 – Effect of age and sex on activity-sociability trait.** Summary of the linear mixed-effect model and null-full model comparison results.

**Supplementary Table 4**

*Model:* Exploration ~ Age + Sex + (1|Group/id)

| **Fixed effects** | **Estimate** | **Std. error** | **z value** | **p-value** |
| --- | --- | --- | --- | --- |
| (Intercept) | 0.435 | 0.271 | 1.602 | 0.109 |
| Age | -0.131 | 0.036 | -3.562 | <0.001 *** |
| Sex (Male) | 0.655 | 0.284 | 2.306 | 0.021 * |

*Comparison with null model:* Likelihood ratio test - χ2 = 21.001, p < 0.001 ***

*(Significance code:* 0.01 ‘*’, 0.000 ‘***’*)*

**Supplementary Table 4 – Effect of age and sex on exploration.** Summary of the linear mixed-effect model output and null-full model comparison results.

**Supplementary Table 5**

| **Name** | **Activity-Sociability** | **Affiliation** | **Exploration** |
| --- | --- | --- | --- |
| **Group 1** | | | |
| Alibi | -1.00 | 0.21 | -0.25 |
| Bowi | 1.76 | -0.87 | -0.87 |
| Dukki | -0.32 | 0.28 | 0.34 |
| Impromptu | 1.70 | 0.64 | 0.64 |
| Mooi | -0.50 | 0.18 | -0.59 |
| Oui | 1.47 | -0.35 | 0.47 |
| Tebbi | -0.37 | 2.54 | 1.14 |
| Toffi | -0.71 | -0.49 | -0.22 |
| Tutudetuu | -0.80 | -2.32 | 1.89 |
| Urbi-et-orbi | -0.83 | 0.46 | -1.16 |
| Walibi | -0.41 | -0.28 | -1.38 |
| **Group 2** | | | |
| Castello | -0.74 | 1.49 | -0.49 |
| Elmo | 0.41 | -0.14 | 1.08 |
| Emerald | 0.28 | -0.04 | 0.80 |
| Etten | -0.19 | -0.21 | -0.55 |
| Freggel | 1.60 | -0.60 | 2.17 |
| Hacienda | -1.21 | -1.10 | -0.82 |
| Lageveen | 0.21 | 0.21 | -0.26 |
| Magelaen | -0.61 | 0.60 | -1.13 |
| Monopoly | 2.62 | -0.83 | 0.70 |
| Nellie | -1.22 | -1.43 | 1.22 |
| Rizzo | -0.11 | -0.11 | -1.23 |
| Rox | -0.26 | 1.18 | -0.27 |
| Saboteur | -0.71 | 0.24 | -0.88 |
| Sjors | 1.48 | -1.06 | 1.74 |
| Soest | 0.04 | 0.24 | -0.96 |
| Stas | -0.79 | 1.09 | -0.65 |
| Tamayo | -0.80 | 0.46 | -0.47 |
| **Group 3** | | | |
| Driel | 0.30 | 0.78 | -0.26 |
| Horsten | 0.12 | -1.91 | -0.81 |
| Kluivers | -1.07 | -0.22 | 1.42 |
| Sollie | 0.66 | 1.35 | -0.35 |

**Supplementary Table 5 – Personality scores of the individuals.** Summary of the identities of the individuals, the groups they belonged to, and personality scores from the three personality dimensions.

**Supplementary Table 6**

*Model:* Problem-focused coping score ~ Activity-sociability + Affiliation + Exploration + Age + Sex + (1|Group)

| **Fixed effects** | **Estimate** | **Std. error** | **t value** | **p-value** |
| --- | --- | --- | --- | --- |
| (Intercept) | 0.644 | 0.184 | 3.499 | 0.006 ** |
| Activity-sociability | -0.219 | 0.133 | -1.639 | 0.135 |
| Affiliation | -0.044 | 0.095 | -0.460 | 0.656 |
| Exploration | 0.087 | 0.098 | 0.887 | 0.398 |
| Age | -0.003 | 0.022 | -0.141 | 0.890 |
| Sex (Male) | 0.077 | 0.227 | 0.339 | 0.742 |

*(Significance code:* 0.001 ‘**’*)*

**Supplementary Table 6 – Effect of personality on problem-focused coping.** Summary of the linear mixed-effect model output.

**Supplementary Table 7**

*Model:* Emotion-focused coping score ~ Affiliation + Exploration + Age + Sex + (1|Group)

| **Fixed effects** | **Estimate** | **Std. error** | **t value** | **p-value** |
| --- | --- | --- | --- | --- |
| (Intercept) | 3.080 | 0.337 | 9.119 | 0.011 * |
| Affiliation | -0.862 | 0.143 | -6.003 | 0.026 * |
| Exploration | 0.411 | 0.193 | 2.122 | 0.167 |
| Age | -0.244 | 0.062 | -3.943 | 0.058 |
| Sex (Male) | -2.261 | 0.331 | -6.818 | 0.020 * |

*(Significance code:* 0.01 ‘*’*)*

**Supplementary Table 7 – Effect of personality on emotion-focused coping.** Summary of the linear mixed-effect model output.

**Supplementary Fig. 4**

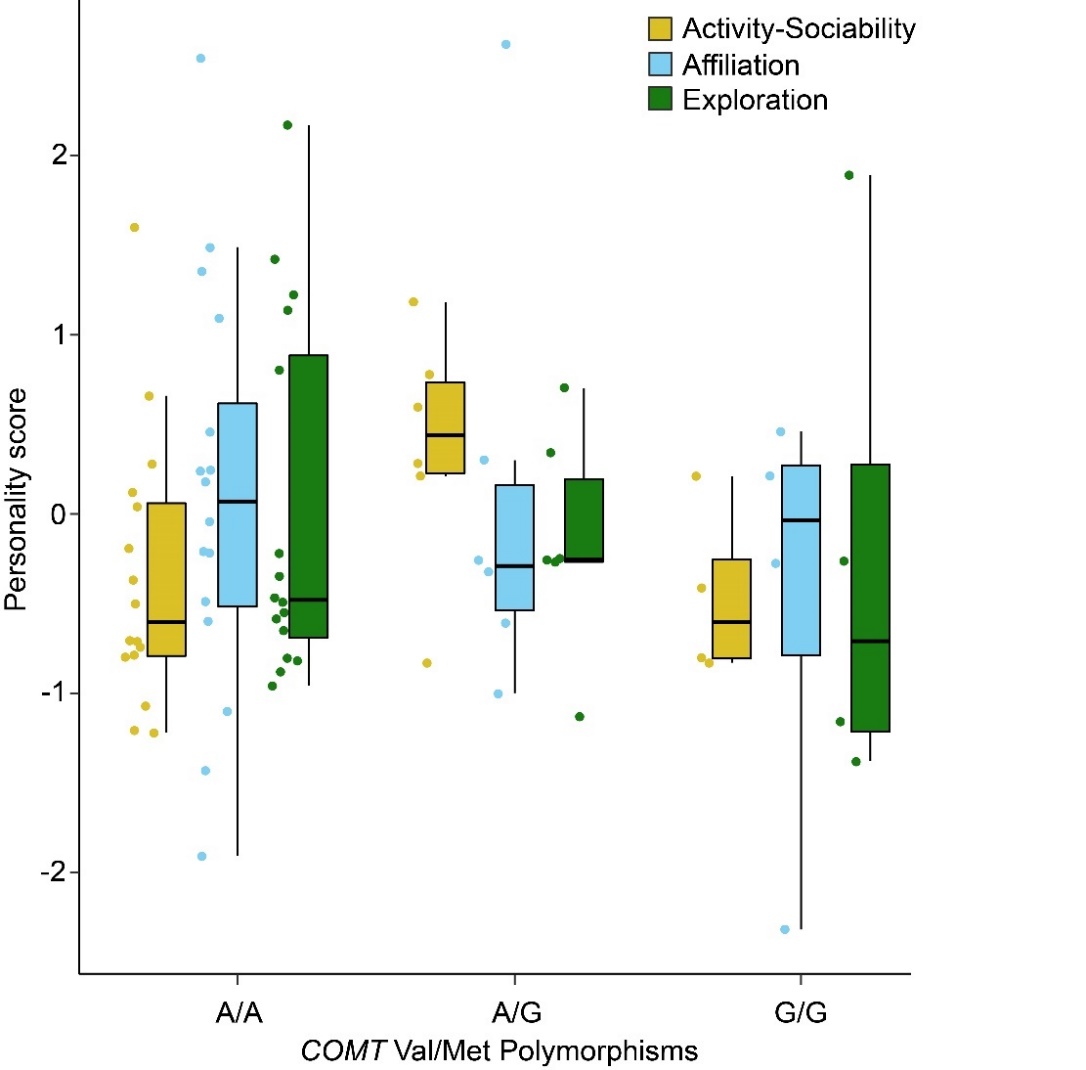

**Supplementary Fig. 4 - *COMT* Val/Met encoding polymorphism and personality traits.** The box plot shows the personality scores from the three traits and their association with *COMT* Val/Met genotype. Individual data points (n=26) are represented using solid dots. Boxes represent interquartile ranges, and whiskers represent the upper and lower limits of the data. The horizontal bars within the boxes represent the median values.

**Supplementary Table 8**

| **Identity and sex of the individuals** | | | **Date of birth** |
| --- | --- | --- | --- |
| **Group 1** | | | |
| Dukki ♀ |  |  | 13/10/2017 |
| Mooi ♀ |  |  | 16/08/2017 |
| Alibi ♀ |  |  | 18/10/2021 |
| Toffi ♀ |  |  | 30/09/2017 |
| Walibi ♀ |  |  | 10/09/2012 |
|  | Bowi ♀ * |  | 18/10/2020 |
| Urbi-et-orbi ♀ |  |  | 13/10/2014 |
|  | Tutudetuu ♂ |  | 11/11/2018 |
|  | Oui ♀ * |  | 18/09/2020 |
| Tebbi ♀ |  |  | 31/08/2017 |
|  | Impromptu ♂ * |  | 29/01/2021 |
| **Group 2** | | | |
| Rizzo ♂ * |  |  | 24/10/2009 |
| Castello ♀ |  |  | 14/01/2008 |
| Saboteur ♀ |  |  | 17/08/2007 |
|  | Tamayo ♀ |  | 20/10/2017 |
| Hacienda ♀ |  |  | 19/01/2008 |
| Magelaen ♀ |  |  | 14/08/2007 |
|  | Soest ♀ |  | 11/05/2015 |
|  |  | Emerald ♀ | 09/04/2019 |
|  |  | Freggel ♂ | 09/04/2020 |
|  |  | Elmo ♂ * | 15/02/2021 |
|  | Etten ♀ |  | 13/08/2016 |
|  | Rox ♀ |  | 18/04/2017 |
|  | Lageveen ♀ |  | 17/10/2018 |
|  | Monopoly ♂ |  | 08/06/2019 |
|  | Stas ♀ |  | 21/08/2014 |
|  |  | Nellie ♀ | 14/06/2019 |
|  |  | Sjors ♂ * | 27/02/2021 |
| **Group 3** | | | |
| Sollie ♂ |  |  | 12/12/2015 |
| Horsten ♂ |  |  | 20/10/2017 |
| Kluivers ♂ |  |  | 22/10/2017 |
| Driel ♂ |  |  | 04/12/2017 |

**Supplementary Table 8 – Details of the monkeys in the study.** The identity, sex, relatedness, and date of birth of the participating macaques from the three groups are summarised. The placement of the macaques from left to right indicates mother-offspring relationships. For example, Bowi is the daughter of Walibi in Gr.1. Individuals who were not included in the *COMT* Val/Met polymorphism analyses are denoted with ‘*’.

**Supplementary Note 3 – Additional details on enclosure and diet of the animals.**

All enclosures had multiple enrichment structures, including slides made from firehoses, plastic toys, wooden structures of various heights, platforms, climbing stairs, a plastic pool (except for Gr.3), and a tree trunk. Note that the exact enrichment materials may have differed to some extent across enclosures. Extra enrichment materials, such as branches and paper containers, were provided when available. The indoor enclosure had concrete floors covered with sawdust bedding, while the outdoor enclosures were covered with natural soil and sand substrates. All indoor enclosures were temperature controlled and maintained a constant temperature of 22˚C. The feeding of the animals always took place in the indoor enclosures. The diet consisted of monkey pellets in the morning, placed in feeding cans attached to the enclosure fences, and vegetables in the afternoon, with an occasional seed mix (corn, sunflower, etc.) being thrown in to stimulate foraging behaviour. Water was available 24/7 ad libitum in both indoor and outdoor enclosures.

**Supplementary Fig. 5**

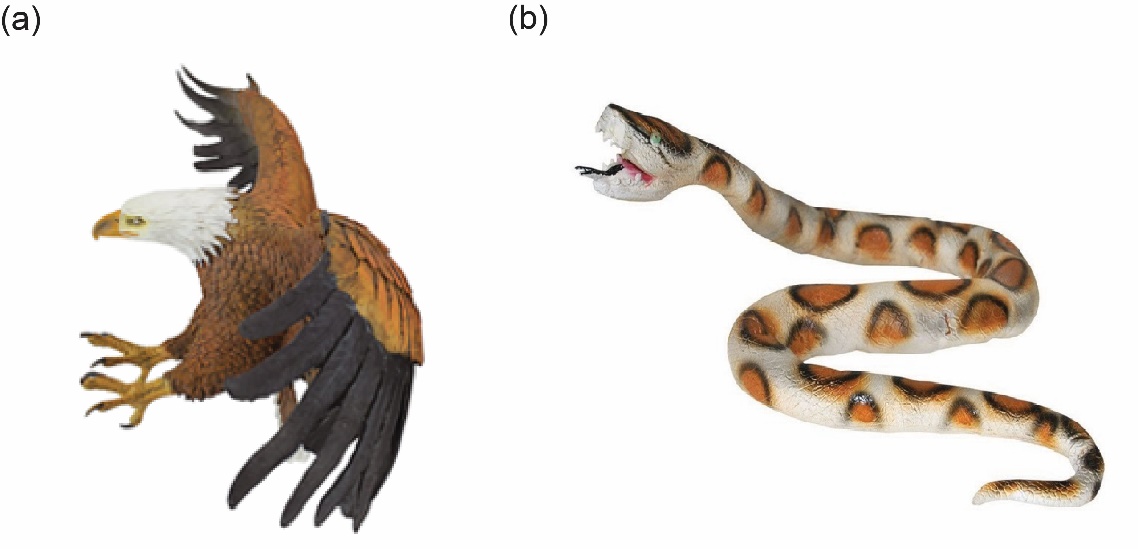

**Supplementary Fig. 5 – Predator models used in the study. (a)** Bird of prey resembling a hawk, (b) Snake resembling a python.

**Supplementary Fig. 6**

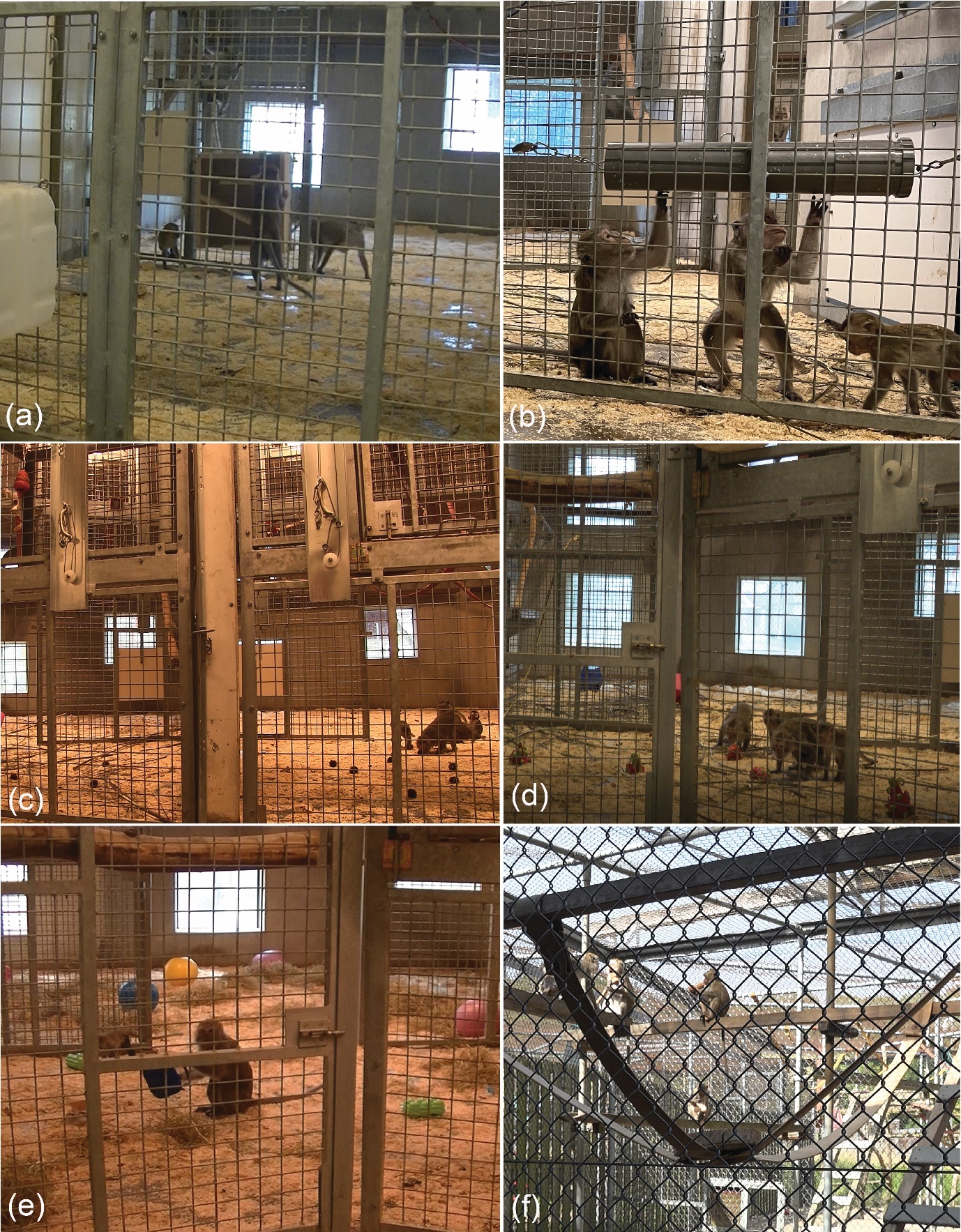

**Supplementary Fig. 6 – Novelty experiments for personality assessment.** (a) Monkeys operating a food puzzle box to retrieve food rewards, (b) Upon rotating a food pipe puzzle, monkeys are obtaining rewards, (c) monkeys inspecting novel food rambutans and (d) dragon fruits, (e) monkeys holding egg containers and (d) massage rollers.

**Supplementary Table 9**

| **Behavioural variable** | **ICC** | **95% CI; lower, upper** | **F-value** | **p-value** |
| --- | --- | --- | --- | --- |
| **Close ground** | **0.397** | 0.049, 0.659 | 2.317 | 0.013 |
| **Far** | **0.799** | 0.620, 0.899 | 8.971 | < 0.001 |
| Locomotion | **0.548** | 0.240, 0.756 | 3.429 | < 0.001 |
| Foraging | **0.319** | -0.040**,** 0.605 | 1.938 | 0.039 |
| **Conspecific aggression** | **0.588** | 0.295**,** 0.780 | 3.859 | < 0.001 |
| **Predator aggression** | **0.544** | 0.234**,** 0.753 | 3.386 | < 0.001 |
| Autogroom | 0.044 | -0.315**,** 0.393 | 1.093 | 0.405 |
| Scratch | 0.042 | -0.317**,** 0.391 | 1.088 | 0.410 |
| Freeze | **0.547** | 0.238**,** 0.755 | 3.415 | < 0.001 |
| Yawn | **0.669** | 0.412**,** 0.827 | 5.053 | < 0.001 |
| **Conspecific affiliation** | **0.540** | 0.229**,** 0.751 | 3.351 | < 0.001 |

**Supplementary Table 9 – Intraclass correlation (ICC) results on coping-related behavioural variables**. ICC values, 95% Confidence interval (CI), F- and p-values are summarised. Repeatable variables following a cutoff of 0.3 are presented in bold fonts. Behavioural variables highlighted in bold fonts were used in the final exploratory factor analysis.

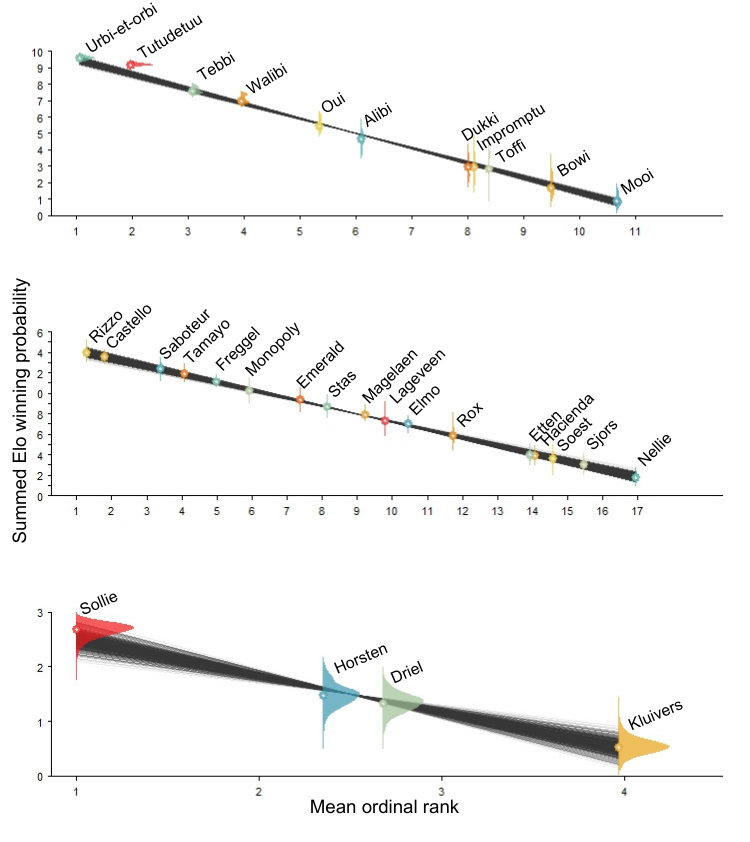
**Supplementary Fig. 7**

**Supplementary Fig. 7 – Dominance hierarchies of the groups.** Based on submissive behaviours, the mean Elo-winning probabilities and 95% CI values of the monkeys (n=32) are plotted against their ordinal ranks. From top to bottom: Gr1. to Gr.3.
